# Cellular dynamics and epidermal specialization during Arabidopsis floral development at single-cell resolution

**DOI:** 10.1101/2025.09.01.672611

**Authors:** Siye Chen, Manuel Neumann, Zhaohui He, Frederic Carew, Cezary Smaczniak, Caroline Braeuning, Sarah Richter, Yingjue Zhang, Xinkai Zhou, Kerstin Kaufmann

**Author notes:** Correspondence should be addressed to Kerstin Kaufmann. These authors contributed equally to this work (Siye Chen, Manuel Neumann).

## Abstract

Cellular specialization underlies the functional diversification of floral organs, yet the regulatory networks driving epidermal differentiation remain poorly understood. Here, we constructed a single-nucleus transcriptomic atlas of developing *Arabidopsis thaliana* flowers by profiling over 70,000 nuclei. For annotation of single-cell identities and differentiation status, we additionally generated tissue- and stage-specific transcriptomic datasets using nine fluorescent marker lines spanning four floral developmental stages. Guided by this integrative approach, we focused specifically on the floral epidermis at single-cell resolution and resolved 22 transcriptionally distinct epidermal cell populations. Gene regulatory network analyses identified key transcription factors essential for epidermal differentiation, highlighting MYB16 as a central regulator within the petal epidermis. Developmental trajectory and chromatin-binding analyses demonstrated that MYB16 dynamically orchestrates gene expression programs involved in epidermal identity establishment, cuticle biosynthesis, and stress resilience. Moreover, we demonstrate MYB16 interacts with DRMY1, a regulator of organ growth robustness. Complementation assays confirmed the non-redundant role of MYB16 in epidermal cell fate specification, distinct from other MYB transcription factors. This study provides a comprehensive cellular and regulatory framework for floral epidermal specialization, advancing our understanding of how temporal transcriptional dynamics integrate with spatial cell fate decisions during flower development.

## Introduction

Flower development serves as a multi-organ model system to understand cell and organ differentiation in plants, facilitated by dynamic changes in gene activities that allow cells to acquire unique identities and functions in reproduction. Thus, cell type– specific transcriptomes are key to uncovering developmental trajectories and cell fate decisions in the flower. While high-resolution data exist for less complex organ systems such as roots and leaf^1–3^, an equivalent resource for the flower remains to be established. Single-cell RNA sequencing (scRNA-seq) enables the identification of cellular differentiation and developmental trajectories at single-cell level. However, the annotation of single-cell omics data is challenged by the complexity of organ-type specific modifications in cell fate in the flower^4^. Early flower development in Arabidopsis starts with the initiation and outgrowth of the floral meristem^5^, associated with molecular patterning processes that determine the positions and identities of floral organs. After around 3 days, the sepal primordia arise, followed by the initiation of petals, stamens and carpels. Growth rates and timing of cellular differentiation patterns are unique to each organ type, and are associated with dynamic changes in metabolic status, such as photosynthetic activity and cell morphology. While the floral meristem activity gets terminated upon carpel initiation, meristematic activity is maintained in specific cell populations, such as the procambium (forming the vasculature) and in the carpel margin meristem (giving rise to ovules) during floral organ differentiation stages. During later stages of organ differentiation, anthers and carpels form specific structures that enable gametogenesis and fertilization. Ovule development commences at stage 9, while the tapetum is formed in anthers at stage 8^6,7^. Around stage 9 of flower development, six nectaries develop as outgrowth of stamen filament-associated epidermal and sub-epidermal layers of the receptacle.

In more general terms, the three major tissue types - epidermis, ground tissue and vasculature - show different degrees of specialization across the floral organs. While epidermal cell structures and cell type composition are largely organ-specific, current knowledge suggests a more conservative vasculature anatomy in the different organ types. Thus, generating tissue- and stage-specific transcriptome data for the Arabidopsis flower is essential for reliable annotation of single cell transcriptomics experiments.

The functional anatomy of epidermal cell structures is remarkably different between floral organ types, associated with specific functions such as visual and chemical pollinator attraction (petals and nectaries), pollination (stigma) and protection (anthers, sepals). A number of transcription factors regulating epidermal differentiation in leaves have been identified. For example, MYB16 and MYB106 are closely related R2R3-MYB transcription factors that regulate the development and patterning of epidermal cells, including cuticle formation and trichome morphogenesis. They are partially redundant and act as positive regulators of epidermal cell outgrowth and wax biosynthesis, with MYB106 also playing a role in suppressing cell elongation in petals and promoting cell flattening. Ectopic MYB16 expression causes abnormal stomatal cluster formation, illustrating the link between stomatal development and cutin synthesis^8^ *MYB16-amiRNA myb106* plants furthermore revealed compromised nanoridge formation in petal conical cells and stamen filaments^9^. However, while they were found to act in concert with the AP2-ERF TF SHINE1 in wax biosynthesis, knowledge on their target gene networks remain limited.

In this project, we established a single cell transcriptomics atlas to comprehensively map cellular differentiation trajectories in the Arabidopsis flower. To facilitate cluster annotation, we generated a resource for stage-specific epidermal, ground tissue and vasculature-associated cells. In a next step, using targeted single cell transcriptomics of the epidermis in the developing flower, we determined distinct sets of transcription factors controlling epidermal cell morphologies in specific floral organs. We further elucidated downstream networks of the master regulatory transcription factor (TF) MYB16 and show that it functions as a temporally dynamic regulator of epidermal differentiation. By integrating single-nucleus RNA sequencing (snRNA-seq), gene regulatory network (GRN) inference, and chromatin immunoprecipitation sequencing (ChIP-seq), we reconstructed MYB16-centered regulatory modules that operate across floral developmental stages. Pseudotime trajectory analysis of epidermal cell populations revealed that MYB16 governs distinct sets of transcriptional targets in early, mid, and late differentiation phases, including genes involved in cuticle formation, cell wall organization, stress response, and developmental timing. Together, our work provides a comprehensive single-cell framework for dissecting epidermal gene regulation in the flower, and positions MYB16 as a central coordinator of temporal patterning, cell identity, and developmental robustness.

Flower development serves as a powerful multi-organ model to understand cell and organ differentiation in plants, driven by dynamic gene expression changes that enable cells to acquire distinct identities and functions essential for reproduction. Cell type–specific transcriptomes are thus crucial to elucidate developmental trajectories and cell fate decisions within flowers. While detailed single-cell data exist for simpler organ systems such as roots, a comparably comprehensive resource for floral tissues remains lacking. Single-cell RNA sequencing (scRNA-seq) has become a valuable method for revealing cellular differentiation and developmental pathways at single-cell resolution. However, annotating single-cell data from flowers is challenging due to the complex, organ-specific differentiation patterns and cell fate dynamics involved.

Early floral development in Arabidopsis begins with floral meristem initiation and outgrowth, coupled with molecular patterning events that define floral organ positions and identities. Sepal primordia emerge around day three, followed sequentially by petals, stamens, and carpels. Each organ type exhibits distinct growth rates and differentiation timings, accompanied by changes in metabolic status, photosynthetic activity, and cell morphology. While floral meristem activity ceases upon carpel initiation, specific meristematic cell populations persist, such as those in the procambium (forming vasculature) and carpel margin meristem (generating ovules). Later differentiation stages involve specialized structures within anthers and carpels that facilitate gametogenesis and fertilization. Ovules and tapetal tissues begin differentiation around floral stages 9 and 8, respectively, while six nectaries form at stage 9 from epidermal and sub-epidermal layers adjacent to stamen filaments.

The three principal floral tissue types—epidermis, ground tissue, and vasculature— display varying degrees of specialization among different floral organs. Epidermal cells exhibit substantial organ-specific diversity, whereas vascular anatomy remains comparatively conserved across organs. Consequently, generating tissue- and stage-specific transcriptomic data for Arabidopsis flowers is critical for accurate annotation and interpretation of single-cell transcriptomic experiments.

Epidermal cell structures in floral organs fulfill specialized functions, including visual and chemical attraction (petals and nectaries), pollination (stigma), and protection (anthers and sepals). Several transcription factors controlling epidermal differentiation in leaves have been identified; for instance, the closely related R2R3-MYB transcription factors MYB16 and MYB106 regulate epidermal cell development, cuticle formation, and trichome morphogenesis. They function partially redundantly as positive regulators of epidermal cell outgrowth and wax biosynthesis. MYB106 additionally suppresses cell elongation in petals, promoting cell flattening.

Misexpression of MYB16 causes aberrant stomatal clustering, highlighting links between stomatal and cuticle development. Studies of MYB16-amiRNA myb106 plants have demonstrated impaired nanoridge formation in petal conical cells and stamen filaments. Despite their recognized roles in wax biosynthesis alongside AP2-ERF TF SHINE1^9^, the gene networks regulated by these transcription factors remain incompletely characterized.

In this study, we present a single-cell transcriptomic atlas to comprehensively map cellular differentiation trajectories in the Arabidopsis flower. To enhance cluster annotation, we established tissue-specific transcriptomic datasets for epidermal, ground tissue, and vasculature-associated cell types at multiple developmental stages. Utilizing targeted single-cell profiling of the floral epidermis, we identified transcription factors governing epidermal cell morphologies across floral organs. We further characterized downstream regulatory networks orchestrated by the transcription factor MYB16, revealing its role as a temporally dynamic regulator of epidermal differentiation. By integrating single-nucleus RNA sequencing (snRNA-seq), gene regulatory network (GRN) inference, and chromatin immunoprecipitation sequencing (ChIP-seq), we reconstructed MYB16-centric regulatory modules active throughout floral development. Pseudotime trajectory analysis showed MYB16 modulates distinct gene sets across early, mid, and late differentiation phases, including genes involved in cuticle formation, cell wall organization, stress responses, and developmental timing. Together, our findings provide a comprehensive single-cell regulatory framework for understanding floral epidermal gene regulation and position MYB16 as a key integrator of temporal patterning, cellular identity, and developmental robustness in floral tissues.

## RESULTS

### A cellular expression atlas of floral organ differentiation

To investigate cellular differentiation pathways in the flower, with a particular focus on the epidermis, we generated a comprehensive single-nucleus RNA sequencing (snRNA-seq) dataset by integrating data from developing inflorescences, mature flowers, and an epidermis-enriched samples (**Fig. 1a**). For the inflorescence samples, data were collected from three different Arabidopsis accessions (*Col-0*, *Bur-0* and *Cvi-0*), and found to be highly reproducible across these accessions (**Supplementary Fig. 1**). The epidermis-enriched dataset was generated using Fluorescence-Activated Nuclei Sorting (FANS) of developing flowers expressing pro*ML1*-driven nuclear localized GFP. Confocal imaging confirmed epidermis-specific GFP expression in the outermost floral cell layers (see also the following section). After processing and quality control, we retained 71,875 high-quality nuclei (median of 884 genes per nucleus), which were clustered into 37 transcriptionally distinct groups (**Supplementary Fig. 1**). To evaluate the composition of each cluster, we assessed the distribution of nuclei from different ecotypes and stages (**Fig. 1b**). Most clusters displayed contributions from multiple samples, indicating that clustering was primarily driven by transcriptional identity rather than sample origin.

**Fig. 1:**
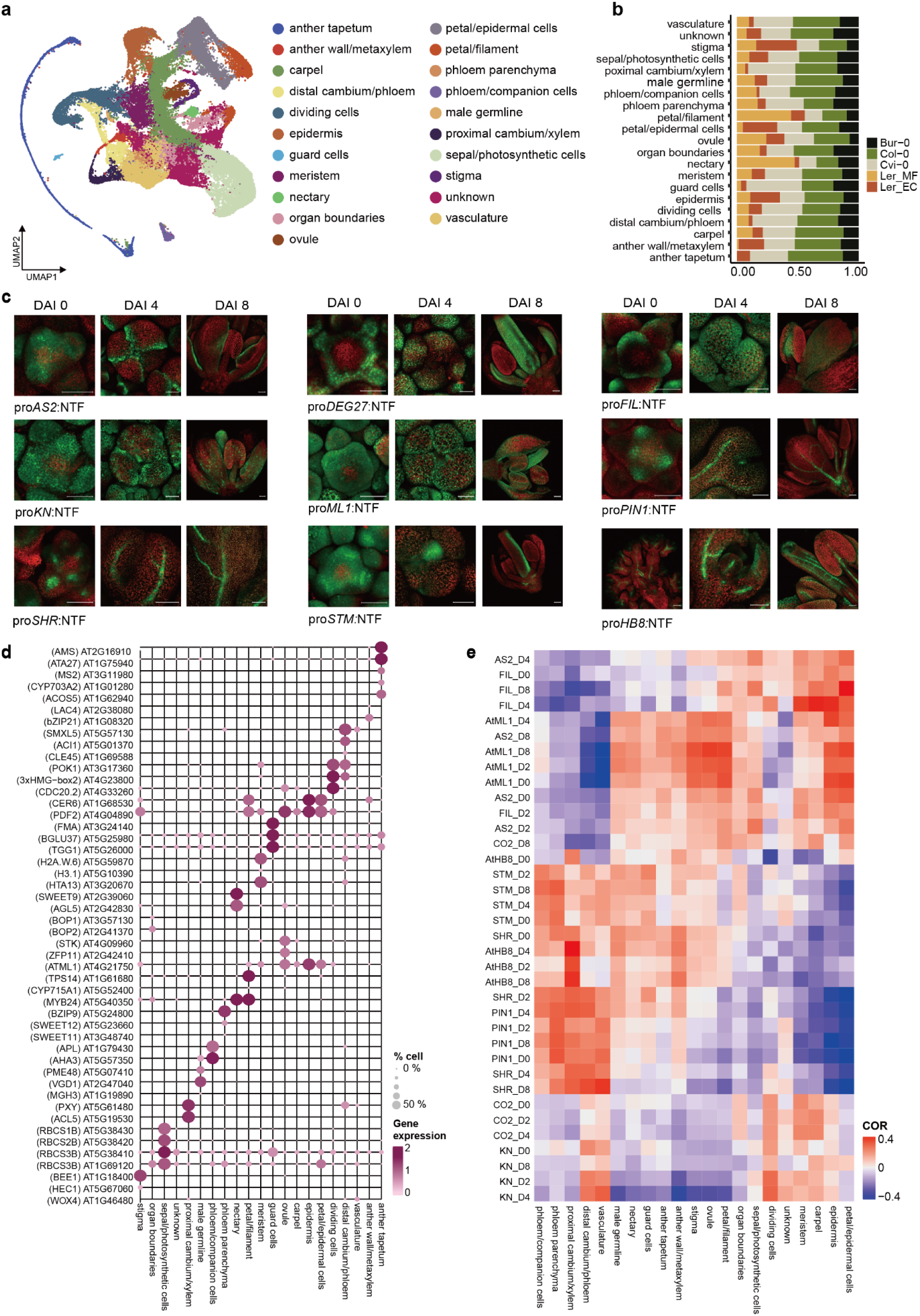
Single-nuclei transcriptomics atlas of Arabidopsis flower development. **a** UMAP embedding of 71,875 nuclei from developing and mature flowers of four Arabidopsis ecotypes, including an epidermis-enriched dataset. Ler_MF, mature flower form Ler; Ler_EC: epidermal cells from Ler. **b** Relative contribution of each input sample to the 21 annotated transcriptional groups. **c** Representative confocal images of floral tissues from nine GFP reporter lines used to define tissue-specific domains for FANS-RNA-seq atlas. Scale bars, 50 μm. DAI, days after induction. **d** Bubble heatmap of known marker gene expression across annotated clusters. Bubble size represents the proportion of expressing nuclei; color indicates average expression. **e** Spearman correlation heatmap between cluster-averaged snRNA-seq profiles and FANS-RNA-seq datasets. COR, Correlation coefficient.

Clusters corresponding to male germline and nectaries were predominantly derived from mature flowers, while the epidermis cluster was enriched in nuclei from the pro*ML1*-FANS dataset. The anther tapetum cluster lacked nuclei from mature flowers, consistent with the known onset of tapetal programmed cell death during late anther development ^10–12^.

The annotation of the snRNA-seq data collection made it necessary to dissect complex patterns of cell differentiation across floral organs and stages/timepoints, We therefore used a combination of available transcriptomics data^13,14^ and newly generated FANS RNA-seq data, in order to distinguish the complex patterns of cell differentiation across different stages and floral organs. FANS RNA-seq data were generated from nuclear membrane-tethered GFP reporter plant lines^15,16^ driven from the promoters of *ARABIDOPSIS THALIANA HOMEOBOX GENE*-8 (*ATHB*-8; cambium^17^), *SHORTROOT* (*SHR*; pericycle^18^), *SHOOTMERISTEMLESS* (*STM*; meristematic stem cells^19,20^), *KNOLLE* (*KN*; division-competent cells^21^), *MERISTEM LAYER1* (*ATML1*; epidermis^22,23^), *FILAMENTOUS FLOWER* (*FIL*; floral organ primordia; at later stages abaxial-shifted^24^); *ASYMMETRIC LEAVES2* (*AS2*; adaxial side of floral organ primordia, later organ margins, anther wall/stomium and inner medial domain in carpel^25,26^); *DEG27* (*CO2*; organ primordia except anther^27^); *PIN-FORMED 1* (*PIN1*; auxin maxima^28^). All constructs were transformed into the system for synchronized floral induction^29^(*proAP1::AP1-GR ap1 cal*) to isolate fluorescently sorted nuclei of specific cell populations enriched in stage 0 (IM), stage 3 (2 days after induction (DAI)), stage 5-6 (4 DAI) and stage 8-10 (8 DAI). All datasets were generated in three biological replicates with high reproducibility (**Supplementary Fig. 2**). Differential expression analysis across timepoints revealed between 2 and 10 significantly regulated genes per tissue (fold change >2, adjusted p < 0.05), with the largest transcriptional shifts observed in *FIL* and *AS2* tissues, while procambium-like populations (e.g., *AtHB8*) remained transcriptionally stable over time (**Supplementary Table. 6**).

Cluster annotation in the combined snRNA-seq dataset was guided by multiple layers of transcriptomic evidence. Expression of known marker genes enabled the identification of both major and rare floral cell types, including phloem (*BZIP9*)^30^, companion cells (*APL*)^30^, proximal and distal cambium (*PXY*, *SMXL5*)^31–33^, stigma (*BEE1*)^34^, ovule (*STK*)^35^, nectary (*SWEET9*)^36^, and male germline (*MGH3*)^37^ (**Fig. 1d**). Notably, epidermal populations exhibited strong organ specificity, while vascular-associated clusters showed organ-independent transcriptional signatures.

To systematically map clusters to spatial domains, we computed Spearman Correlations between cluster-averaged snRNA-seq profiles and FANS-RNA-seq tissue references (**Fig. 1e**). This analysis revealed strong concordance between epidermal clusters and *AtML1* samples, vascular clusters and *SHR*, or *ATHB8* samples, as well as dividing cell clusters and the *KN*-enriched dataset. These results highlight the resolving power of our FANS-based reference atlas in capturing both broad tissue domains and heterogeneous, transient populations such as meristematic and proliferative cells. Its integration with snRNA-seq data enabled spatially grounded annotation, even in the absence of uniquely defining marker genes. For clusters lacking organ-specific markers, we further incorporated bulk transcriptomic data from the TraVaDB floral atlas, which provides organ- and stage-resolved expression profiles. This comparison enabled confident annotation of transcriptionally ambiguous clusters, such as sepal and carpel (**Supplementary Fig. 1**). In order to facilitate data exploration and reuse by the broader research community, we developed an interactive web portal (Integrated atlas:https://www.epiplant.hu-berlin.de/cellxgene/view/atha_flower_integrated_atlas.h5ad/; epidermis-enrich atlas: https://www.epiplant.hu-berlin.de/cellxgene/view/atha_flower_epidermis_atlas.h5ad/) using the CellxGene platform^38,39^. Users can visualize gene expression across annotated clusters, compare marker specificity across tissues and stages, and interactively explore the floral transcriptional landscape.

Altogether, we present a spatially and temporally resolved single-nucleus transcriptomic atlas of Arabidopsis floral tissues, constructed through the integration of snRNA-seq, tissue-specific FANS-RNA-seq, and public reference datasets. By capturing the transcriptional identity of both abundant and rare floral cell types, this atlas provides a high-resolution framework for exploring gene regulatory programs, cellular differentiation, and tissue patterning in flowers. As an open-access resource, it offers a powerful foundation for dissecting plant organogenesis at single-cell resolution.

### Epidermis-specific regulatory networks uncover candidate hub transcription factors in floral organs

To explore transcriptional regulation within the floral epidermis, we first isolated epidermal nuclei from developing Arabidopsis flowers using FANS based on pro*AtML1*::GFP expression, which marks the outermost epidermal layers (**Fig 2c and Supplementary Fig 3**). The sorted nuclei were subjected to snRNA-seq and subsequently integrated with our broader floral cell atlas. Unsupervised clustering of the epidermis-enriched dataset resolved 22 transcriptionally distinct epidermal cell populations (**Fig. 2a**). To assign spatial identity to each cluster, we selected top marker genes and compared their expression profiles to stage- and organ-resolved reference datasets from the TRAVA database. Clusters 0 and 1 were assigned to sepals, cluster 6 to ovules, cluster 10 to filaments, cluster 11 to stigmatic tissue, and cluster 21 to anthers. Notably, four petal-associated clusters (5, 4, 16, and 18) exhibited stage-specific gene expression signatures: cluster 5 was enriched for genes expressed at floral stage 5, while clusters 16, 4, and 18 aligned with stages 4, 2, and 1, respectively (**Supplementary Fig. 3**). Based on these signatures, we annotated cluster 5 as representing young petal epidermis.

**Fig. 2:**
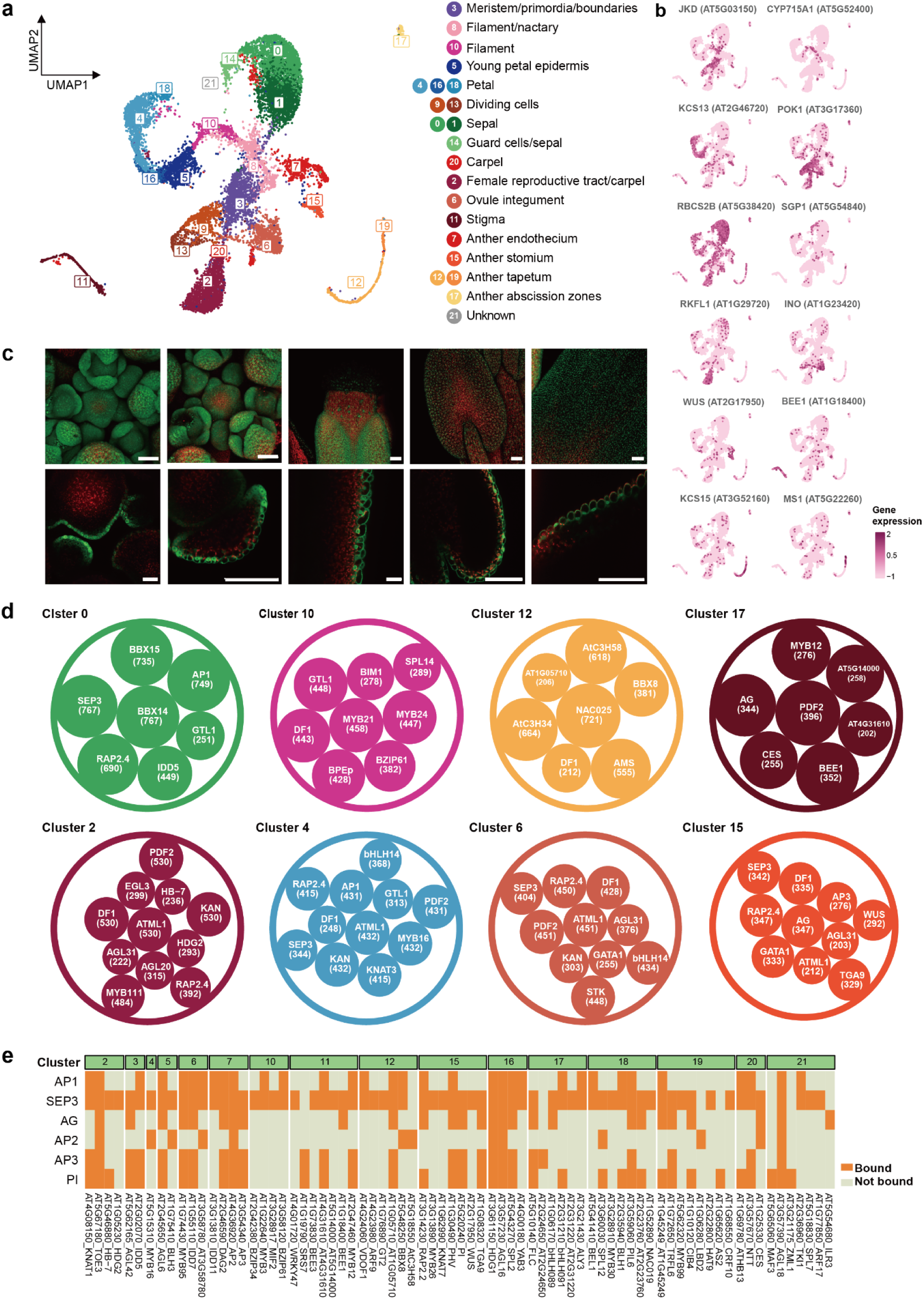
Integration of single-cell nuclear transcriptomics and gene regulatory analysis reveal floral organ-specific epidermal regulatory hubs. **a** UMAP of single-cell RNA-seq data from Arabidopsis epidermal nuclei **b** Expression patterns of selected known marker genes projected onto the UMAP **c** Representative confocal images showing epidermis-specific GFP signals in different floral organs. Scale bars, 50 µm. **d** Representative hub transcription factors (frequency >200 times) identified from GRN analysis for selected clusters. The number of predicted downstream targets was indicated. **e** Heatmap showing binding of floral homeotic TFs to hub genes across clusters, based on ChIP-seq peak overlap.

These annotations were further validated by the expression of well-studied marker genes. For example, *RUBISCO SMALL SUBUNIT 2B* (*RBCS2B*; sepal)^40^, *INNER NO OUTER* (*INO*; ovule)^41^, and *BRENHANCED EXPRESSION1* (*BEE1;* stigma)^34^ showed cluster-specific enrichment consistent with prior knowledge. We also identified specialized epidermal cell types, including meristematic/boundary cells (*JACKDAW* (*JKD*))^42^, anther tapetum (*MALE STERILITY1* (*MS1)*, *ASD*)^43,44^, anther stomium (*WUSCHEL* (*WUS*))^45^, anther stomium (*MYB DOMAIN PROTEIN26* (*MYB26*))^46^, *NAC SECONDARY WALL THICKENING PROMOTING FACTOR1* (*NST1*)^47,48^, further supporting the robustness of our annotation strategy **(Fig. 2c and Supplementary 2**).

To systematically uncover regulatory dynamics across epidermal cells, we reconstructed cluster-specific gene regulatory networks (GRNs) using the GENIE3 algorithm^44^, focusing on regulatory interactions between known transcription factors (TFs) and all expressed genes. For each epidermal cell group, the top 5,000 regulator-target interactions were selected based on interaction weight as defined by feature importance scores. Within each group, TFs were then ranked by the numbers of predicted downstream targets. To increase stringency, we retained only those TF–group combinations in which the TF exhibited more than 50 predicted unique target genes. This final high-confidence set of hub regulators was considered as candidate master transcriptional coordinators within each epidermal subtype (**Fig. 2d and Supplementary Table 1**).

Across multiple representative clusters, hub TFs showed strong concordance with organ identity and the biological roles of their predicted targets. For instance, in the sepal clusters (0 and 1), hubs such as *SEPALLATA3* (*SEP3*), and *APETALA1* (*AP1*) were prominent. The involvement of *SEP3* and *AP1*—two floral homeotic genes— suggests that their regulatory influence extends beyond early meristem patterning into organ-specific epidermal specification (**Fig 2d**). In the ovule integument cluster (6), *SEEDSTICK* (*STK)* was identified as a key hub, in line with its role in ovule identity and integument differentiation^49^ (**Fig 2d**). In the anther stomium (cluster 15), *WUS* was a central hub TF. Distinct from its stem cell regulatory role in the shoot apical meristem, *WUS* in the anther regulates stomium and septum cell fate (**Fig 2d**). Loss-of-function mutants fail to trigger programmed cell death in these cells, blocking anther dehiscence and pollen release, highlighting a unique regulatory pathway for anther opening^45^. In the anther tapetum clusters (12 and 19), *ABORTED MICROSPORES* (*AMS*) functioned as the dominant hub, consistent with its well-established role in tapetum development and pollen wall formation^50^. In the stigma cluster (cluster 11), *AGAMOUS* (*AG*) was identified as a hub transcription factor (**Supplementary Table 1**). Although *AG* is primarily known for its early role in specifying stamen and carpel identity, *in situ* hybridization studies have shown that its expression becomes spatially restricted at later stages of flower development.

Specifically, *AG* RNA accumulates in differentiated gynoecial cell types, including the stigmatic papillae, suggesting that *AG* may contribute to stigma maturation and cellular specialization beyond its canonical role in organ identity establishment^51^.

More prominently, *BR Enhanced Expression 1* (*BEE1*) stood out as a central hub TF in this cluster (**Supplementary Table 1)**. The brassinosteroid-responsive bHLH transcription factor *BEE1* is specifically expressed in the stigma and stylar transmitting tract, where it acts redundantly with *HALF-FILLED* (*HAF*) and *BR ENHANCED EXPRESSION 3* (*BEE3*) to regulate reproductive processes. These include extracellular matrix deposition to support pollen tube passage and the onset of stigmatic cell death following pollination^34^. The identification of *BEE1* as a hub highlights the importance of brassinosteroid-mediated transcriptional regulation in establishing stigma receptivity and enabling successful fertilization. In the filament cluster (10), *AUXIN RESPONSE FACTOR8* (*ARF8*) emerged as a prominent hub transcription factor (**Fig. 2**). Although it had a relatively modest number of predicted targets, several were functionally relevant, including the *FRAGILE FIBER1* (*FRA1)*, encoding a Kinesin-like protein with an N-terminal microtubule binding motor domain that mediates cellulose microfibril orientation (**Supplementary Table 1**). *ARF8* is a key component of the auxin signaling pathway and has been previously implicated in filament elongation, likely by modulating genes involved in cell wall remodeling and mechanical reinforcement^52^. Its identification as a hub in this cluster supports a role for auxin-responsive transcriptional regulation in driving filament growth and structural integrity.

Among the identified TFs, MYB family members are prominent for their regulatory specificity and diversity. For example, *MYB16* was mainly enriched in petal clusters (cluster 4, 16, 18 and 5), regulating *ECERIFERUM 3* (*CER3)* and *SHINE1* (*SHN1)*, both involved in cuticle biosynthesis (**Fig. 2d and Supplementary Table 1**). *MYB26* was dominant TFs in anther stomium (15), while *MYB24*, in the filament cluster (10), targeted *PIN-FORMED 3* (*PIN3,* auxin efflux) and *TERPENE SYNTHASE14* (*TPS14*), a gene expressed in filament according to GUS reporter assays, suggesting a role in filament cell development (**Fig. 2d**)^53^. In the female reproductive tract/carpel cluster (2) and petal cluster (16 and 5), *MYB111* targeted *ATP-BINDING CASSETTE12* (*ABCG12*) and *HYDROPEROXIDE LYASE 1* (*HPL1*), implicating it in cuticular lipids transportation and defense-related metabolism^54,55^. *MYB124* (*FOUR LIPS (FLP*)) exhibited broader activity, acting as a hub across multiple clusters including dividing cells (clusters 9, 13), carpel (2), and anther-related clusters (17 and 19) (**Fig. 2d, Supplementary Table 1**). This pattern is in line with a role in cell cycle regulation during differentiation^56^. Indeed, *MYB124* was shown to be broadly expressed in early developing flowers, and controls ovule initiation at later stages in development^57^. Similarly, *MYB21* emerged as a recurrent hub across filament, stigma, petal, and carpel clusters, consistent with its known regulation by jasmonate signaling during floral maturation^58^.

In addition to well-characterized epidermal regulators, we identified several novel cluster-specific hub transcription factors that may contribute to previously unrecognized regulatory modules in floral epidermal development. *CCCH-TYPE ZINC FINGER PROTEIN34* (*AtC3H34*) emerged as a prominent hub in the anther tapetum (cluster 12). Its predicted targets include *AUXIN RESPONSE FACTOR 9* (*ARF9*), *DICER-LIKE3* (*DCL3)*, and *3-KETOACYL-COA SYNTHASE 21* (*KCS21*), implicating it in auxin signaling, chromatin remodeling, and cuticular lipid biosynthesis, indicating that *AtC3H43* may integrate hormonal and chromatin-mediated programs to support lipid-mediated tapetum function (**Fig. 2d, Supplementary Table 1**). In the stigma-associated cluster (cluster 11), *WRINKLED4 (WRI4)*, a transcription factor known for its role in lipid metabolism, was identified as a major hub. WRI4 was predicted to target key cuticle-related genes such as *LONG-CHAIN ACYL-COENZYME A SYNTHETASE1* (*LACS1*), *3-KETOACYL-COA SYNTHASE 20* (*KCS20*), and *ABCG12*, along with hormone-responsive genes including SNF1-RELATED PROTEIN KINASE 2.5 (*SNRK2.5*), and *BEE1*, suggesting its involvement in coordinating hormonal signaling with barrier lipid deposition during stigma development (**Supplementary Table 2**). In the filament cluster (cluster 10), *BZIP61* was highlighted as a tissue-specific hub regulator. Its targets span cell wall biosynthetic enzymes (*CELLULOSE SYNTHASE A1* (*CESA1*), *FASCICLIN-LIKE ARABINOGALACTAN PROTEIN18 (FLA18)*, *CER3*), hormone-related factors (*AUXIN RESPONSE FACTOR 16* (*ARF16*), *TRICHOME CELL SHAPE 1*(*TCS1*)), and chromatin modifiers (*SET-DOMAIN GROUP2 (SDG2)*), pointing to a multifaceted role in regulating filament elongation and epidermal integrity (**Fig. 2d, supplementary table 1 and 2**). These findings expand the landscape of floral epidermal transcriptional control by revealing tissue-specific TFs that integrate hormonal, metabolic, and epigenetic inputs.

In conclusion, our analysis reveals a complex yet organized transcriptional landscape within the floral epidermis. By combining high-resolution single-nucleus transcriptomics with gene regulatory network inference, we predicted both known and novel transcription factors that likely coordinate the spatial specialization of epidermal cell types. The discovery of hub regulators with organ-specific activity underscores the modularity of floral epidermal regulation, while the identification of previously uncharacterized TFs expands the repertoire of candidate regulators.

These findings provide a mechanistic framework for understanding how transcriptional programs shape floral surface identity and offer a valuable resource for future functional studies into epidermal patterning during reproductive development.

#### Homeotic transcription factors target hub genes associated with different organ epidermal development

To investigate how floral organ identity transcription factors, regulate epidermal cell differentiation, we integrated the GRN analysis with ChIP-seq datasets for key homeotic MADS-box proteins, including *AP1*, *AP2*, *AP3*, *PI*, *AG*, and *SEP3*^59–62^. By mapping homeotic TF binding profiles to cluster-specific hub genes identified from our single-cell transcriptomic atlas, we assessed the extent to which floral regulators directly control organ-specific epidermal transcriptional modules. Overall, we found that around 44% of the 168 hub genes predicted by GRN analysis are bound by at least one homeotic TF. Most predominantly, SEP3 binding was identified for 67 hub genes (∼39%).

In clusters annotated as carpel —including the carpel wall (cluster 20), stigma (cluster 11), ovule integument (cluster 6), and the female reproductive tract (cluster 2)—hub genes such as *CES*, *BEE1*, *IDD5*, *MYB95* and *TOE*3 were found to be bound by *SEP3* and *AG*. These targets are associated with brassinosteroid biosynthesis and integument differentiation, indicating that *AG*–*SEP3* complexes drive a specialized transcriptional program in the carpel epidermis (**Fig. 2e**). In petal-associated clusters (4, 5, 16, and 18), several hub transcription factors—including *PIL6*, *AGL16*, and *HDG1*—were bound by combinations of *AP1*, *AP3*, *PI* and *SEP3*, with *HDG1* and *AGL16* additionally targeted by *AP2*. *MYB16* (cluster 4), a well-characterized regulator of cuticle biosynthesis, was directly bound by *SEP3* and *AP2*, aligning with its established role in defining petal epidermal differentiation.

*PIL6* (hub regulator in cluster 18), which encodes a Myc-related bHLH transcription factor from the PIF3 family, was bound by the petal identity tetramer (*AP3*-*PI*-*AP1*-*SEP3*) **(Fig. 2e)**. It is defined as a component of the phytochrome-mediated red-light signaling pathway, modulating photomorphogenic responses. Floral organ growth is known to be regulated by red light signaling^63^, thus likely via the PHYB-PIF signalling module. Accordingly, *PIL6* may link red-light perception and circadian signaling with petal growth downstream of homeotic TFs.

Together, these results demonstrate that distinct combinations of floral homeotic transcription factors directly regulate functionally specialized epidermal hub genes in a floral organ-specific manner. This reveals a hierarchical regulatory architecture in which MADS-box protein complexes interface with tissue-specific GRNs to orchestrate epidermal differentiation in coordination with floral organ identity. Such coordinated regulation ensures that epidermal features—such as cuticle formation, pollination, and environmental sensing—are precisely aligned with the developmental and functional needs of each organ.

### Temporal dynamics of MYB16-associated gene expression along petal epidermal differentiation trajectories

To investigate the temporal regulatory dynamics underlying epidermal differentiation, we firstly applied RNA velocity analysis to our epidermis-specific snRNA-seq dataset to infer the directionality of cellular differentiation. The velocity vectors indicated a clear trajectory from meristematic to differentiated epidermal cell states, allowing us to define the root and terminus of the pseudotime trajectory (**Supplementary Fig. 4**). Based on this information, we constructed a pseudotime trajectory by integrating cells from petal and meristem clusters (**Fig. 3a**). The trajectory revealed a continuum from meristematic to differentiated epidermal cell states. To validate the biological relevance of this trajectory, we compared it to stage-specific transcriptomic profiles from the TRAVA database, which showed high concordance with our single-cell– derived trajectory **(Fig 3b)**.

**Fig. 3:**
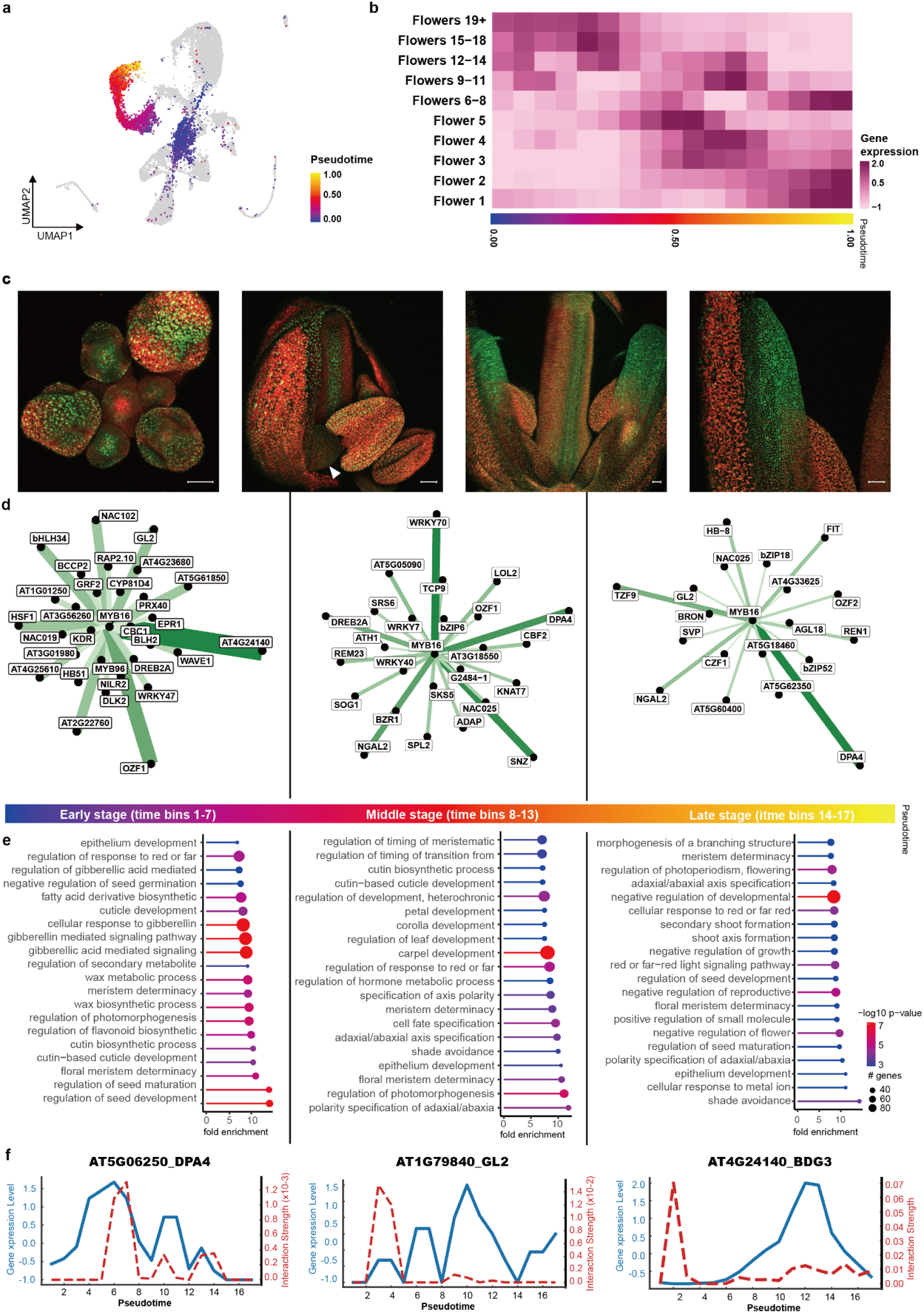
Temporal dynamics of MYB16 activity along petal epidermal differentiation trajectories. **a** Pseudotime trajectory of petal epidermal differentiation reconstructed from epidermis-enriched snRNA-seq data. **b** Comparison of pseudotime stages with TRAVA flower stage–specific transcriptomes. **c** Confocal images showing MYB16-GFP signals in different floral organs at different stages. Arrowhead indicates petal primordia; Scale bars, 50 µm. **d** Interactions dynamics of representative MYB16 target genes along pseudotime. Developmental time bins grouped into early (1–7), mid (8–13), and late (14–17) phases. **e** GO enrichment analysis of MYB16 targets at each developmental stage. **f** Stage-specific regulation of MYB16 target transcription factors across petal epidermal differentiation.

Focusing on MYB16, a hub transcription factor identified in petal clusters through our earlier GRN analysis, we analyzed the expression dynamics of its predicted target genes along the pseudotime trajectory. These genes displayed distinct temporal expression patterns, indicative of a coordinated regulatory program operating during petal epidermal differentiation. To better resolve these dynamics, we divided the pseudotime trajectory into three transcriptionally distinct phases—early (time bins 1– 7), mid (8–13), and late (14–17)—based on comparisons between the transcriptome profiles of individual pseudotime bins and stage-specific datasets from the TRAVA database (**Fig. 3b** and **3d**). This approach allowed us to capture gene expression programs associated with MYB16 activity, helping to resolve the temporal progression of petal epidermal differentiation.

To enable identification of potential direct target genes of MYB16 by ChIP–seq, we generated transgenic lines expressing a MYB16–GFP fusion protein driven by the native MYB16 promoter. To examine the expression pattern of the fusion protein, we first examined MYB16–GFP expression by confocal microscopy. MYB16–GFP was strongly expressed in petal epidermal cells and localized predominantly to the nucleus, consistent with its function as a transcription factor (**Fig. 3c**). At early stages, MYB16–GFP expression was broadly detected in the epidermal cells of sepals, carpels, and petal primordia, whereas during later stages (stage 11–12), expression became enriched in petal epidermal cells, highlighting a dynamic regulatory role during petal development.

Having validated appropriate expression and nuclear localization of the fusion protein, we next performed ChIP–seq using developing flowers from MYB16–GFP plants. Two independent biological replicates were generated using anti-GFP antibody immunoprecipitation. Both replicates showed robust enrichment, although the number of peaks identified in each replicate was variable. To improve the stringency and interpretability of the data, we merged the read count data from both biological replicates and conducted subsequent peak calling on the combined dataset. This analysis identified 3,248 significant MYB16 binding sites (log2 fold change > 2.0, FDR < 0.05). A gene was defined as a potential MYB16 target if a binding site was located within 3 kb upstream to 1 kb downstream of its transcription start site (TSS), resulting in 4,151 potential target genes. Similar to other transcription factors, MYB16 binding events were predominantly enriched in promoter-proximal regions (**Supplementary Fig. 5**). To link MYB16 binding to dynamic gene regulation during petal development, we compared GRN-predicted targets with MYB16-bound genes identified in the ChIP–seq analysis. This comparison revealed 351 overlapping genes, representing 52.24% of all predicted targets (Fig. 6; Supplementary Table 2). We visualized the expression dynamics of these high-confidence MYB16 targets along the petal pseudotime trajectory to investigate their regulatory patterns across developmental phases (**Fig. 3d**).

To elucidate how MYB16 directs petal epidermis differentiation over time, we analyzed its transcription factor targets phase by phase. In the early phase, several of these TFs are closely associated with epidermal cell fate determination and cuticle formation. For example, *GLABRA2 (GL2)*, a well-established regulator of trichome initiation and epidermal identity, was among the early *MYB16* targets, suggesting a direct transcriptional link between MYB16 and terminal differentiation pathways^64^.

*HSF1* (AT4G17750), a heat shock transcription factor, was also bound by *MYB16*, indicating a potential role in maintaining transcriptional stability or stress resilience during the early phases of cell fate commitment^65^. Among the targets in the early phase, *BODYGUARD3* (*BDG3*; AT4G24140) stood out due to its strong regulatory weight in the network (**Fig 3c**). *BDG3* is associated with cuticle development and epidermal barrier formation; although its function is less well characterized than its homolog *BDG1*, its expression pattern and predicted regulatory connectivity suggest a role in reinforcing surface integrity^65–67^. The strong inferred interaction between *MYB16* and *BDG3* implies that *MYB16* may promote epidermal differentiation not only via classical transcriptional regulators but also through activation of enzymes for cuticle maturation. Given that *bdg* mutants typically exhibit increased cuticle permeability and organ fusion phenotypes, MYB16-driven expression of *BDG3* during early developmental windows may be critical for establishing an effective epidermal barrier^66^. Additionally, *MYB16* targets several transcription factors involved in hormonal and stress responses—including *WRKY47* and *DEHYDRATION-RESPONSIVE ELEMENT BINDING PROTEIN 2* (*DREB2A*)—highlighting a broader role in buffering environmental cues to ensure robust execution of the epidermal gene expression program^68^. Notably, *WRKY47* also functions as a hub transcription factor in cluster 11 (stigma) (**Supplementary Table 1**), suggesting that *MYB16* may influence stigma development by integrating epidermal differentiation with stress-responsive regulatory networks in reproductive tissues.

At mid developmental stages, *MYB16* continued to perform extensive transcriptional control over a distinct set of TFs. Several of these mid-phase targets are involved in stress adaptation, developmental timing, and chromatin regulation, suggesting a shift in MYB16’s function towards maintaining cellular differentiation and limiting transcriptional plasticity. Notably, *NGATHA-LIKE PROTEIN3* (*NGA3*/*DPA4),* a negative regulator of cell proliferation during petal development^69^, was among the top-ranked mid-stage targets. Its suppression by *MYB16* likely serves to prevent reactivation of stem cells–like transcriptional programs and to enforce commitment to epidermal differentiation^70^. Similarly, *DREB2A*, a key mediator of drought and osmotic stress responses, remained a persistent target from early to mid-stages, highlighting the continuous need to suppress immune- or stress-responsive transcriptomic shifts that could disrupt cuticle development^68^.MYB16 also targeted *WRKY DNA-BINDING PROTEIN 70* (*WRKY70*), which involved in salicylic acid signaling and defense regulation, reinforcing its role in shielding developmental progression from stress-induced perturbations^71^. In parallel, MYB16 bound several TF genes involved in structural organization and spatial patterning, including *KNOTTED-LIKE HOMEOBOX OF ARABIDOPSIS THALIANA 7* (*KNAT7*), which is implicated in secondary cell wall biosynthesis - but so far not in epidermis differentiation^72^.This interaction points to a possible role for MYB16 in regulating the cell wall architecture of the epidermal layer.

At late stages of epidermal development, MYB16 retained control over a specific cohort of transcription factors, reflecting its role in stabilizing terminal cell identity and actively suppressing residual transcriptional plasticity. Persistent interaction of *DPA4*, in particular, highlights MYB16’s continued role in safeguarding against ectopic activation of stem cell–associated programs. In addition to these enduring targets, MYB16 regulated several late-specific repressors and chromatin-related TFs, including *SHORT VEGETATIVE PHASE* (*SVP*) and *AGAMOUS-LIKE 18* (*AGL18*), both MADS-box transcription factors involved in flowering time control and growth arrest^73–75^. Their MYB16-mediated regulation may serve to reinforce terminal differentiation. Interestingly, late-stage MYB16 targets also included *FIT*, a central regulator of iron homeostasis, and *GL2*, a marker of epidermal identity. The regulation of *GL2* across both early and late pseudotime phases suggests that MYB16 contributes to sustaining epidermal gene expression well beyond initial fate commitment.

To further assess the biological relevance of this stage-specific regulatory cascade, we performed Gene Ontology (GO) enrichment analysis along the pseudotime axis. For each time bin, we selected the top 100 genes based on interaction weight and identified enriched functional categories associated with their expression. This analysis revealed a temporal progression of biological processes: early pseudotime bins were enriched for terms such as meristem determinacy, epithelium development, and gibberellin mediated signaling pathway; mid-stage bins showed enrichment for regulation of photomorphogenesis, adaxial/abaxial axis specification, and cutin biosynthetic process; while late-stage bins prominently featured categories such as shade avoidance, and negative regulation of growth (**Fig. 3e and Supplementary fig 5**). These results reinforce our proposed model that MYB16 functions as a temporal master regulator, coordinating epidermal cell fate specification, structural maturation, and transcriptional stabilization throughout petal development. Together, these analyses highlight the role of MYB16 as a temporal master regulator guiding sequential stages of petal epidermal differentiation.

#### Stage-specific regulation of representative MYB16 target genes

To further dissect how MYB16 dynamically regulates its target genes during petal development, we integrated pseudotime-resolved gene expression profiles with predicted MYB16 binding strength. By combining transcriptional dynamics with regulatory interaction intensity, we aimed to reveal how MYB16 modulates key developmental programs across distinct stages. Representative target genes displayed diverse expression–binding relationships, illustrating stage-specific mechanisms through which MYB16 coordinates petal epidermal differentiation and maturation (**Supplementary fig 6**).

*DPA4* and *BDG3* (AT4G24140) were predicted by GRN analysis, directly bound by MYB16 in ChIP-seq, and significantly up- (*DPA4*) vs. downregulated (*BDG3*) in the *myb16* and *myb16 myb106* mutants, identifying them as high-confidence direct targets of MYB16 during petal development. *DPA4* exhibited multiple transient expression peaks along pseudotime, each closely followed by a surge in MYB16 binding (**Fig. 3f**). This inverse relationship suggests that MYB16 acts to suppress *DPA4* reactivation at specific developmental windows, thereby contributing to the stabilization of petal cell identity. Given that *DPA4* promotes stem cell niche formation by repressing *CUC* boundary genes, timely inhibition by MYB16 likely prevents reversion to meristematic programs during petal differentiation ^70,76^. *BDG3*, which is involved in cuticle biosynthesis and epidermal integrity, showed strong MYB16 binding at early stages, although its expression remained low at this time. At later stages, *BDG3* expression markedly increased, suggesting that MYB16 promotes *BDG3* activation through an indirect or multi-step regulatory pathway, thereby fine-tuning the timing of cuticle deposition during epidermal maturation (**Fig. 3f**).

*GL2*, another predicted target and ChIP-bound gene, was also differentially expressed (P-adj < 0.05), although the fold change was modest (log2FC > 0). Transient peaks of MYB16 binding preceded declines in GL2 expression, suggesting that MYB16 modulates *GL2* activity to coordinate epidermal lipid metabolism and cell identity. The modest change in *GL2* expression may reflect its role as a core epidermal regulator whose expression is finely balanced rather than strongly switched during petal differentiation (**Fig. 3f**).

Together, these findings highlight the stage-specific and multifunctional nature of MYB16-mediated transcriptional control. By acting as both a repressor and activator in a context-dependent manner—and targeting genes involved in developmental progression, immune regulation, and environmental response—MYB16 emerges as a central coordinator of epidermal cell fate.

### MYB16-centered gene regulatory networks and transcriptional control

MYB16, the closest homolog of MYB106, has been implicated in epidermal differentiation, with prior studies suggesting functional redundancy between the two transcription factors^9^. To further dissect their roles, we generated novel stable *myb16* and *myb106* loss-of-function alleles via Cas9-mediated mutagenesis. Compared to wild type, *myb16* mutants exhibited significantly reduced petal, sepal, and carpel lengths (p < 0.01), while *myb106* mutants showed a mild but significant increase in stamen length. In the *myb16 myb106* double mutant, petal and sepal lengths remained significantly altered. These organ-level phenotypes were accompanied by marked changes in epidermal features (**Supplementary Fig. 8**). Light microscopy revealed that double mutant petals appeared more translucent and transmitted more light (**Supplementary Fig. 8**), suggesting defects in cuticular integrity. Consistently, scanning electron microscopy showed that while wild-type petals formed characteristic nanoridges, these structures were entirely absent in the double mutant. This loss of nanoridges corroborates the established role of MYB106 in cuticle patterning and underscores a redundant contribution of MYB16 and MYB106 to the specialized surface architecture of petal epidermal cells **(Supplementary Fig. 8**).

To assess the molecular consequences of MYB16 and MYB106 disruption, we performed RNA-seq on developing flowers from each single mutant and the double mutant (three biological replicates per genotype; **Supplementary Fig. 8**). Differential gene expression analysis revealed that the *myb16 myb106* double mutant exhibited the most extensive transcriptional reprogramming, with 239 unique DEGs not detected in either single mutant (**Fig. 4a**). Moreover, 130 genes were shared between *myb16* and *myb106*, 45 between *myb16* and the double mutant, and 80 between *myb106* and the double mutant. Importantly, 136 genes were consistently misregulated across all three genotypes, representing a core set of targets jointly regulated by MYB16 and MYB106. These findings highlight their partially redundant regulatory roles, with MYB16 exerting a broader impact on gene expression and the double mutant exhibiting evidence of strong functional compensation.

**Fig. 4:**
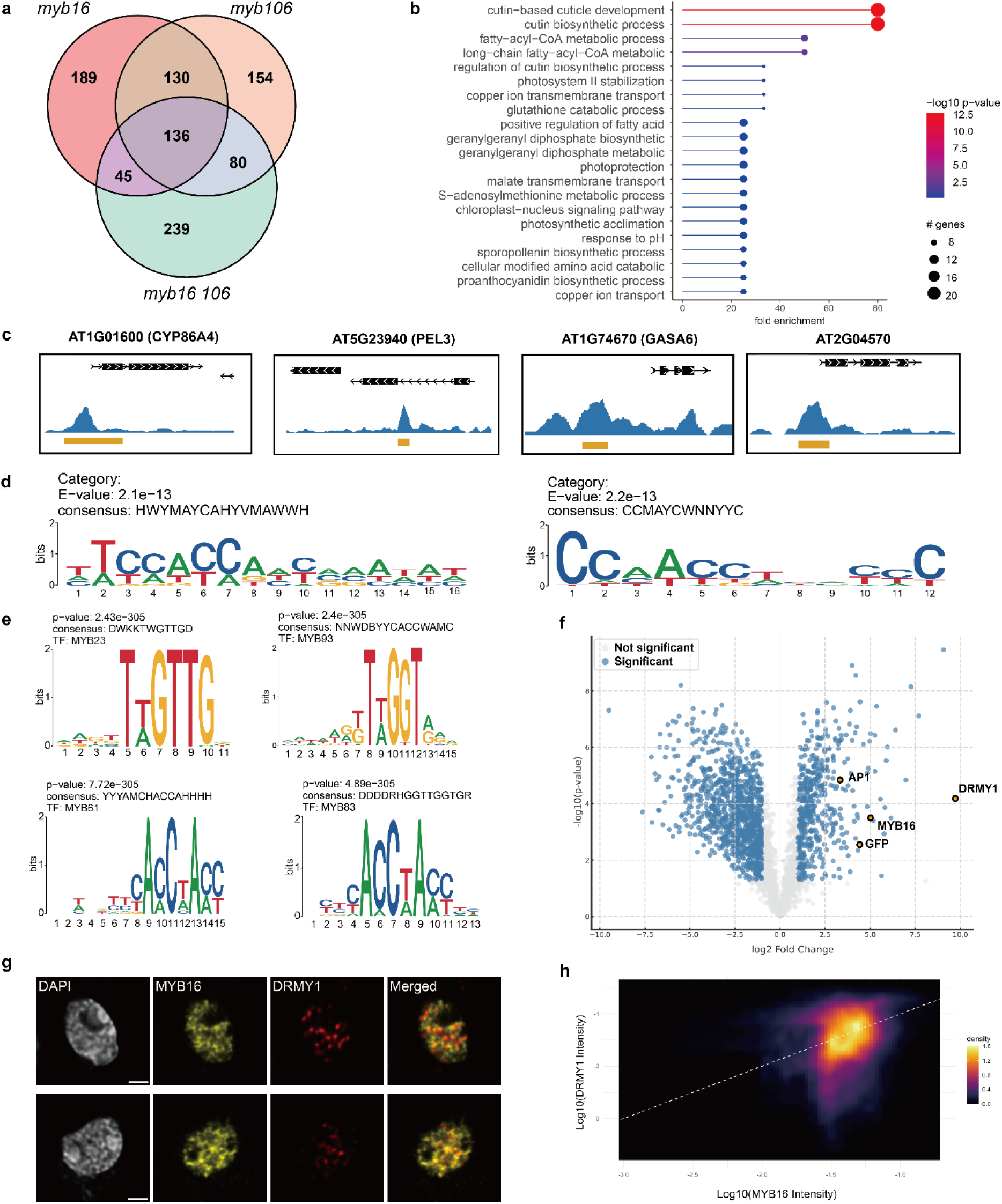
MYB16 directly regulates gene expression and physically interacts with DRMY1 to coordinate epidermis development. **a** Venn diagram showing overlap of differentially expressed genes (DEGs) in *myb16*, *myb106*, and *myb16 myb106* mutants. **b** Gene ontology (GO) enrichment analysis of genes bound by MYB16 (ChIP-seq) and transcriptionally affected in *myb16* and *myb16 myb106* mutants. Dot size indicates the number of genes; color represents –log10 *P* value. **c** Representative MYB16–GFP ChIP–seq peaks in developing floral tissue, highlighting binding events near selected target genes. **d** *De novo* motif discovery from MYB16 ChIP-seq peak sequences using MEME. The top two enriched motifs include a distinct MYB16-specific binding pattern (left) and a canonical MYB-binding element (right). **e** Motif similarity analysis of MYB16 ChIP–seq peak sequences with known MYB family binding motifs. **f** Volcano plot showing enrichment of MYB16-associated proteins identified by immunoprecipitation followed by mass spectrometry (IP–MS). Representative enriched proteins are labeled, including DRMY1. **G** Multiplexed cryoIF images images of Arabidopsis sepal epidermis nuclei co-expressing MYB16–GFP and DRMY1– mCherry. Strong nuclear co-localization is observed. Scale bars, 2 μm. **h** Pixel-by-pixel density scatter plot showing signal correlation between MYB16–GFP and DRMY1–mCherry across optical nuclear slices. Dashed line indicates linear regression fit; the color scale represents pixel density.

To further dissect how these transcriptional changes are directly mediated, we integrated MYB16 binding profiles with differential expression data by intersecting ChIP–seq targets with DEGs from the *myb16* and *myb16 myb106* mutants, yielding 140 candidate direct targets of MYB16.GO enrichment analysis of these genes revealed significant enrichment for biological processes associated with epidermal differentiation and environmental responses (**Fig. 4b**). Representative ChIP–seq peaks at key target genes further support MYB16’s central role in cuticle formation. Strong binding and differential expression were observed for genes involved in cuticle biosynthesis and lipid metabolism including *CYP77A6* (AT3G10570)^77^, *OSP1* (AT2G04570, GDSL lipase)^78^, and *CYP86A4* (AT1G01600) (**Fig. 4c**)^79^. MYB16 also directly regulates (*PERMEABLE LEAVES3* (AT5G23940, *PEL3/DCR*)), a gene required for cuticular nanoridge formation and epidermal impermeability, and AT1G21540 (*AAE9*), encoding an acyl-activating enzyme involved in lipid precursor activation^80,81^. These findings reinforce MYB16’s pivotal role in promoting cuticle integrity and epidermal differentiation. Beyond these classical targets, MYB16 also binds genes associated with cell elongation and growth regulation. For instance, *GASA6* (AT1G74670) are involved in gibberellin signaling and cell elongation, while *MSRB6* (AT4G04830) encodes a methionine sulfoxide reductase that may influence cell wall redox status^82,83^. Additionally, *PRP4* (AT4G38770) encodes a proline-rich protein that may contribute to the mechanical properties of the epidermal cell wall, further implicating MYB16 in morphological control beyond lipid-based processes^84^. (**Supplementary table 2**). Moreover, poorly characterized targets such as *PBL28* (AT1G24030), consistently repressed across all mutant backgrounds, may represent novel components of the MYB16-governed epidermal regulatory network.

To further characterize MYB16’s binding preferences, we performed de novo motif discovery on ChIP–seq peaks using MEME. This analysis identified several enriched sequence motifs, prominently featuring a canonical MYB-binding element (CCMAYCWNNYYC) and an extended motif (HWYMAYCAHYVMAWWH), the latter potentially representing a MYB16-specific binding signature (**Fig. 4d**, **Supplementary Fig. 8**). Additional motifs included sequences with partial MYB-like features (KRANCMRAYGRD) and more complex or poly-purine-rich patterns, which may reflect cofactor binding or chromatin accessibility biases (**Supplementary Fig. 8**). Comparison of MYB16-derived motifs with known MYB transcription factor binding motifs revealed strong similarity to those of MYB23, MYB93, MYB61, and MYB83 (Fig. 4e). Several identified consensus motifs, including TGGTG (MYB23) and CACCWACC (MYB61), match canonical MYB-binding elements characteristic of R2R3-MYB transcription factors (**Fig. 3e**)^85^ Notably, MYB61 and MYB83 are also functionally associated with stomatal aperture and secondary cell wall biosynthesis respectively^86,87^, further supporting the biological relevance of the identified MYB16 binding sites.

### MYB16 interacts with DRMY1, a regulator of organ growth robustness

To elucidate the regulatory mechanisms through which MYB16 governs epidermal differentiation in floral organs, we performed immunoprecipitation–mass spectrometry (IP–MS) using a MYB16–GFP fusion protein. Among significantly enriched proteins, DEVELOPMENT RELATED MYB-LIKE1 (DRMY1, AT3G15030) emerged as a top interactor, showing strong enrichment and statistical significance (**Fig. 4e, supplementary table 4**), indicating physical association with MYB16 *in vivo*.

We next assessed the nuclear colocalization of MYB16 and DRMY1 with multiplex cryo immunofluorescence (multiplex cryoIF) by expressing MYB16–GFP and DRMY1–mCherry fusion proteins in Arabidopsis. Confocal acquisitions revealed colocalization of the two proteins in sepal epidermis nuclei (**Fig. 4g, supplementary Fig. 10**). Quantitative image analysis revealed an asymmetric overlap: ∼75% of nuclear DRMY1 signal colocalized with MYB16, whereas only ∼28% of MYB16 signal overlapped with DRMY1 (Fig. 4h and Supplementary Table 3). This pattern suggests that DRMY1 is largely recruited to MYB16-associated chromatin foci, while MYB16 exhibits broader nuclear distribution. Given that DRMY1 also regulates processes such as cell expansion and hormone responses^88^, its broader nuclear localization likely reflects multifunctional roles. In contrast, MYB16 appears to recruit DRMY1 to specific genomic locations to fine-tune its activity during epidermal differentiation in the sepal.

Supporting this model, comparative transcriptome analysis of *myb16* and *drmy1* mutant flowers (*drmy1* dataset from Wu, et al. ^88^) identified 47 overlapping differentially expressed genes, including genes involved in cuticle biosynthesis, lipid metabolism, and epidermal identity (**Supplementary Table 4**). Notably, genes such as *AT5G52310* (lipid transfer protein) and *AT4G04830* (GDSL lipase) were consistently downregulated in both mutants, suggesting that *MYB16* and *DRMY1* co-regulate a shared transcriptional module.

Together, these results support a model in which MYB16 serves as a core epidermal regulator that conditionally associates with DRMY1 to modulate gene expression in the sepal epidermis. The asymmetric colocalization pattern reflects a modular regulatory mechanism, wherein DRMY1 functions both independently and in cooperation with MYB16 to coordinate epidermis development in different floral organs.

#### Divergent roles of MYB family TFs in petal epidermal differentiation

Building on the expression profiles and gene regulatory network (GRN) analyses derived from our epidermis single-cell transcriptomic dataset, we observed that MYB16 is predominantly expressed in the petal epidermal cluster, whereas MYB24 and MYB26 exhibit specific expression in filament and anther stomium clusters, respectively (**Fig. 5a**). Importantly, *MYB24* and *MYB26* were identified as key hub transcription factors within their respective clusters, suggesting central roles in regulating filament- and anther-specific developmental programs (**Fig. 2d and Supplementary Table 1**). To further investigate potential functional redundancy among these homologous MYB transcription factors, we performed complementation analysis in the *myb16 myb106* double mutant background.

**Fig. 5:**
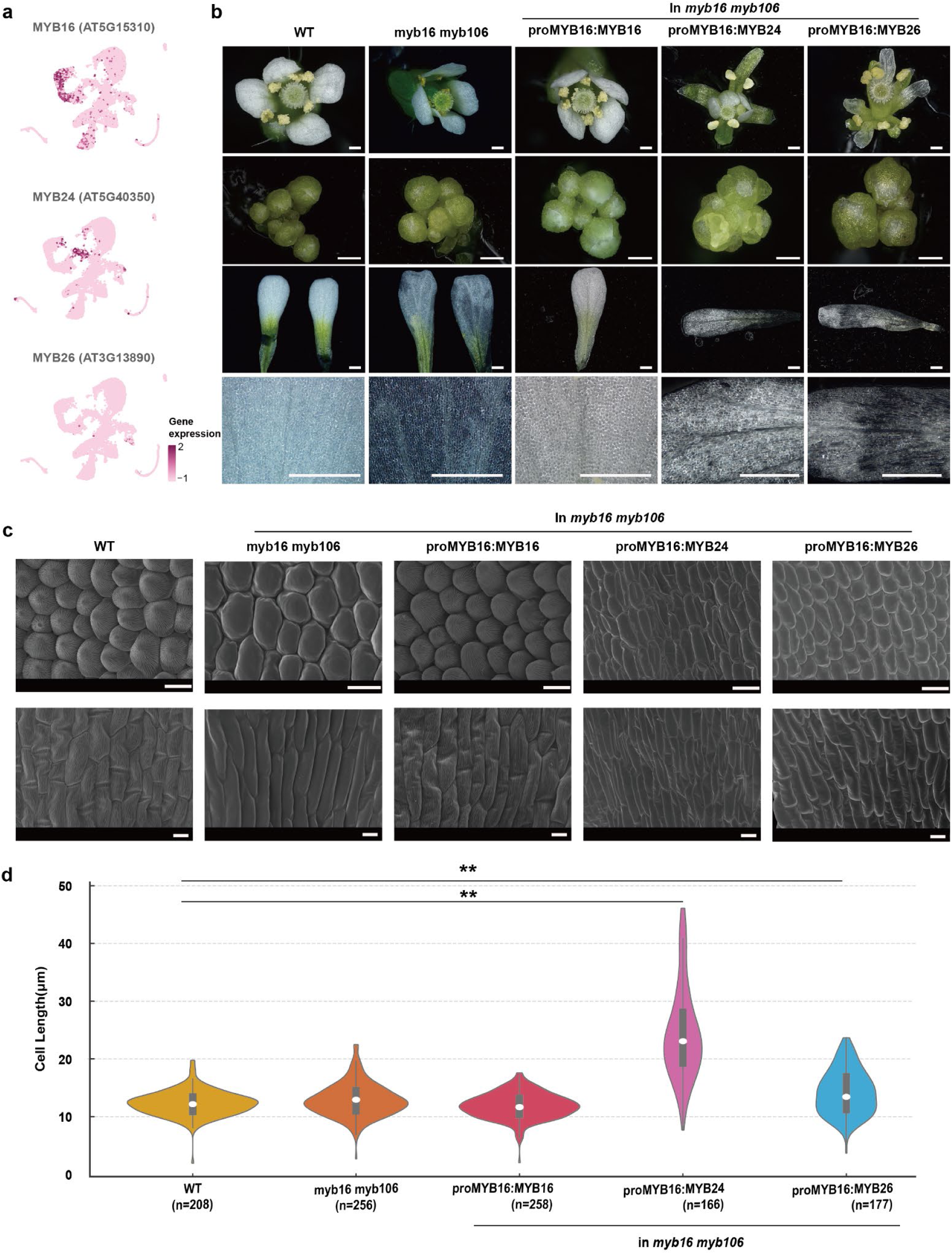
MYB16 is functionally distinct from MYB24 and MYB26 in regulating petal epidermal differentiation. **a** Expression patterns of *MYB16*, *MYB24*, and *MYB26* projected onto the UMAP of floral epidermis single-nucleus transcriptomic atlas. **b** Complementation analysis in the *myb16 myb106* double mutant background using transgenic lines expressing *MYB16*, *MYB24*, or *MYB26* under the *MYB16* promoter. From top to bottom: whole flowers, dissected floral buds, petals, and adaxial petal epidermis under bright-field microscopy. Scale bar, 200μm. **c** Scanning electron microscopy images of upper (top row) and lower (bottom row) part of adaxial petal epidermis. Scale bar, 10 μm. **d** Violin plots showing distribution of adaxial petal epidermal cell length across genotypes. n indicates the number of cells measured. Asterisks indicate significance as determined by Mann–Whitney U test with FDR correction.

Given the severe epidermal defects in *myb16 myb106* mutants (**Fig. 5b)**—including impaired conical cell morphology, loss of nanoridge structures, and increased petal transparency due to cuticle defects—we tested whether *MYB16* homologs could rescue the phenotype by expressing *MYB16*, *MYB24*, or *MYB26* under the control of the native MYB16 promoter.

As expected, MYB16 expression fully restored the petal epidermal phenotype, including the formation of conical epidermal cells, re-establishment of nanoridge structures, and restoration of cuticle integrity, as demonstrated by both optical and scanning electron microscopy (**Fig. 5b and c**). In contrast, neither MYB24 nor MYB26 was able to complement the epidermal defects. Optical microscopy and SEM further revealed that only MYB16 expression reinstated the densely packed, conical epidermal cells characteristic of wild-type petals, whereas MYB24- and MYB26-expressing lines displayed disorganized, elongated epidermal cells with smooth surfaces lacking nanoridges and light-scattering properties (**Fig. 5b and c**).

SEM analysis also revealed pronounced longitudinal expansion of epidermal cells in the MYB24- and MYB26-expressing lines (**Fig. 5c**). Quantitative measurements confirmed that cell length was significantly increased in these lines compared to wild-type and MYB16-complemented plants (**Fig. 5d**). Specifically, Mann–Whitney U tests revealed highly significant differences between WT and MYB24 (FDR = 8.84 × 10⁻⁵³) and WT and MYB26 (FDR = 1.02 × 10⁻⁶). While the FDR value for the MYB16-complemented group (0.051) was marginally above the conventional threshold, these plants displayed cell lengths highly similar to wild type, indicating substantial phenotypic rescue (**Fig. 5d**). The elongation phenotype was particularly pronounced in the petal blade of MYB24- and MYB26-expressing lines, suggesting a shift toward directional cell expansion that replaces the normal conical cell shape in the absence of functional MYB16.

The inability of MYB24 and MYB26 to restore petal epidermal morphology likely reflects their specialized roles in other floral tissues and the distinct regulatory contexts in which they operate. MYB24 functions primarily in stamen development, where it integrates gibberellin and jasmonic acid signaling to promote filament elongation. It forms a transcriptional complex with MYC-type bHLH proteins and is tightly regulated by DELLA and JAZ repressors at both transcriptional and post-translational levels^89^. By contrast, MYB26 plays a central role in initiating secondary wall thickening in the anther endothecium through direct activation of *NST1* and *NST2*. Although MYB26 transcripts are detected in multiple floral organs, its protein accumulation is largely restricted to the endothecium via post-transcriptional regulation, and it fails to function in tissues lacking its NAC-domain co-regulators^46^.

Together, these findings suggest that neither MYB24 nor MYB26 possesses the appropriate spatial context or transcriptional network to drive petal epidermal differentiation, underscoring the tissue-specific functional specialization of MYB16. Future studies employing ChIP-seq, DAP-seq, or transcriptomic profiling in floral tissues will be instrumental in identifying the direct targets of MYB24 and MYB26, and in elucidating the distinct regulatory circuits that underlie their roles in defining tissue-specific cellular growth patterns.

## DISCUSSION

Floral epidermal tissues display striking structural and functional differences, but the gene networks that control their development are still not fully understood. In this study, we built a high-resolution single-nucleus transcriptomic atlas of Arabidopsis thaliana flowers and identified MYB16 as a key transcriptional regulator of petal epidermal differentiation. By integrating snRNA-seq, gene regulatory network analysis, ChIP–seq, and functional experiments, we show that MYB16 shapes epidermal identity by activating and repressing distinct gene sets at different developmental stages.

Our data reveal that MYB16 acts in a temporally dynamic and context-specific manner across different floral organ types. Using the petal as a model system, we found that MYB16 activates genes involved in epidermal identity and cuticle formation. Later, it represses genes associated with stem cell programs and stress responses, ensuring proper maturation of epidermal cells. This phase-specific activity suggests that MYB16 helps to stabilize development by both promoting identity and suppressing unwanted transcriptional noise. Notably, MYB16 functions as both an activator and a repressor, likely depending on cofactor availability and chromatin state.

We also identify DRMY1 as interaction partner of MYB16. Although DRMY1 is involved in broader processes such as ribosome biogenesis and hormone signaling, its specific overlap with MYB16 suggests that it may be recruited to fine-tune epidermal gene activities. The identification of 47 shared differentially expressed genes in *myb16* and *drmy1* mutants—including genes involved in cuticle formation and lipid metabolism—supports this co-regulatory model.

Despite being closely related, MYB24 and MYB26 were unable to replace MYB16 when expressed under the MYB16 promoter in the myb16 myb106 mutant background. These results emphasize the non-redundant, tissue-specific function of MYB16 in the petal epidermis. The inability of these homologs to rescue the phenotype highlights how even closely related transcription factors can have specialized functions depending on tissue context, interaction partners, and regulatory environments.

Our study also demonstrates the value of combining single-cell data with tissue-specific references to resolve the complexity of floral development. The atlas we generated enables high-resolution analysis of cell-type–specific regulatory networks and can serve as a resource for further studies in different genetic backgrounds, species, or environmental conditions.

Nonetheless, there are limitations to consider. While we inferred MYB16 target genes using GRNs and validated many through ChIP–seq, direct time-course data on MYB16 binding across floral stages are lacking. Future studies using time-resolved CUT&Tag or ChIP–seq will help define dynamic binding landscapes.

Moreover, identifying additional cofactors or chromatin remodelers that work with MYB16 and DRMY1 could reveal more about how epidermal fate is regulated in space and time.

In conclusion, this work provides new insight into how transcriptional programs guide floral surface differentiation, positioning MYB16 as a central regulator that links organ identity, epidermal structure, and transcriptional stability. Our atlas offers a valuable framework for future efforts to understand how gene regulation controls organ-specific development at single-cell resolution.

## METHODS

### Plant material and tissue collection

The Columbia (Col-0), Landsberg (Ler), Cape Verde islands (Cvi-0) and Burren (Bur-0) ecotypes of Arabidopsis thaliana were used in this study. Plants were cultivated in a blend of soil and vermiculite, maintaining a 2:1 ratio. The growth conditions are long-day cycles (16 hours of light and 8 hours of darkness), with humidity set at 60%. For the floral inducible plants (*ap1 cal* AP1-GR plants), the plants were induced by applying DEX-induction solution (5μl 20Μm Dexamethasone and 8μl Silwet L-77 in 50 ml distilled water) to the main inflorescence after they bolted and reached 2 cm high. The inflorescences were collected and sap-frozen in liquid nitrogen on day 2, day 4, and day 8 after DEX-induction. The day 0 plants were collected directly when they bolted and reached 2 cm. For the other plants, the inflorescence was collected on day 8 after bolting. The samples for snRNA-seq, RNA-seq, and IP-MS were harvested in liquid nitrogen. Samples designated for ChIP-seq were initially kept on ice before being fixed with 1% formaldehyde. Each type of plant tissue was collected in three biological replicates, allowing for comprehensive analysis and validation of the experimental results.

### Transgenic line generation

Tissue-specific reporter vectors for the genes CO2, KN, SHR, and PIN1 were obtained from NASC (https://arabidopsis.info/, NASC ID: N2106365). The promoters of the genes AS2, FIL, AtHB8, AtML1, and STM were amplified from the Arabidopsis thaliana genome using primers listed in Supplementary Table 9. These promoters sequence subsequently cloned into the pCR8 vector through enzyme digestion, A-tailing, and ligation. The resulting pCR8 vectors containing tissue-specific promoters were then recombined into the pK7WG-INTACT-AT destination vector via an LR reaction to generate GFP transcriptional reporter constructs. The pK7WG-INTACT-AT vector was obtained from VIB-Plant Systems Biology^15^. The pCR8 vector containing an XcmI restriction site was modified from the pCR8/GW/TOPO TA Cloning Kit (Thermo Fisher Scientific, Cat No.: K250020).

To generate the proMYB16:gMYB16-GFP constructs, the proMYB16:gMYB16 (6073 bp) and fragments were amplified and cloned into the pCR8 entry vector, followed by recombination into the pMDC107 destination vector, ultimately generating the pFG1691 vector. The myb16 and myb106 mutants were created using the CRISPR/Cas9 system following the protocol of Schumacher, et al. ^90^, with guide RNA sequences provided in Supplemental Table 9. Genotyping confirmed the mutation sites, and the myb16 line 1-38 and myb106 line 15-5 were selecteFwud for RNA-Seq analysis. The myb16/myb106 double mutant was generated by crossing the two single mutants.

For the promoter swap experiment, the pFG1691 vector was digested with AscI and XbaI, and the CDS of MYB24 and MYB26 was amplified from cDNA. The proMYB16+5′UTR sequence was cloned from the pCR8 proMYB16:gMYB16 construct. These fragments were assembled using the In-Fusion Cloning Kit (Takara Bio USA, Inc.). The myb16/myb106 double mutant was transformed with proMYB16:gMYB16-GFP, proMYB16:MYB24-CDS-GFP, and proMYB16:MYB26-CDS-GFP.

For the colocalization experiment, the proDRMY1:gDRMY1 fragment was amplified from genomic DNA, while the mCherry-containing destination vector backbone was derived from pFG1829. These fragments were assembled using the In-Fusion Cloning Kit and transformed into the Col-0 plant harboring the proMYB16:gMYB16-GFP construct.

All primers used in this study are listed in Supplemental Table 9, and all Agrobacterium-mediated transformations were performed using the floral dip method^91^.

### Confocal microscopy

To investigate the dynamic expression pattern of GFP signals at the reporter line in different developmental stages (D0, D2, D4, and D8 after DEX induction), a Zeiss 800 laser-scanning confocal microscope equipped with 20x objectives and 40 objectives was employed. The GFP signal was excited using a 488 nm laser line, while chlorophyll autofluorescence was excited with a 655 nm laser line to provide a contrasting background delineating the tissue structures. Z-stack images were acquired to capture the three-dimensional structure of the tissues. The subsequent processing of these Z-stack images was carried out with Zeiss imaging software version 3.5 (blue edition).

### Nuclei preparation

The nuclei isolation procedure was based on the protocol by Moreno-Romero, et al.^92^ in Nature Protocols. For the isolation procedure, tissue is first gently crushed using a pestle to break down large chunks, then transferred to a GentleMACS M tube (Miltenyi Biotec) with liquid nitrogen, allowing the nitrogen to evaporate completely. Following this, 5 ml of the Honda buffer (2.5% Ficoll 400, 5% dextran, 0.4M sucrose, 25mM Tris-HCL pH=7.4, 10mM MgCl2) is added to the tube and briefly shaken to prevent freezing. Tissue homogenization is performed using the GentleMACS dissociator (Miltenyi Biotec) at 4 degrees, after which the homogenate is filtered through a 75 µm cell strainer, and the filter is washed with an additional 5 ml of Honda buffer. The filtrate is then centrifuged for 6 minutes at 1000g and 4°C, the supernatant removed leaving behind 1 ml of Honda buffer for nuclei resuspension and filtered again through a 35 µm cap mesh.

For FANS, the sample is divided into two fractions: approximately 100 µl for an unstained control and around 800 µl for staining with 5 µl of 100 mg/ml DAPI, plus 13 µl RiboLock RNase Inhibitor (Thermo Fisher Scientific, Cat No.: EO0382). The sorting tube is prepared with 15 µl of 4% BSA in PBS and 6 µl of RiboLock RNase Inhibitor in a 1.5 ml Eppendorf tube. The FANC is performed by using BD FACSAria Fusion (BD Biosciences). The experiment starts with tubes for a DAPI-unstained control and a non-GFP-stained control. After running the control sample, the sorting sample is recorded and sorted for DAPI signal with GFP-positive nuclei.

### FANS RNA-seq

Total RNA for FANS RNA-seq was isolated using miRNeasy Micro Kit (Qiagen, Cat. No.: 217084). After sorting, nuclei are centrifuged at 1500g for 10 minutes at 4°C, the supernatant mostly removed, leaving about 20 µl, and the nuclei lysed with 350 µl RLT buffer, vortexed for 1 minute. The homogeneous cell lysate is frozen in liquid nitrogen and stored at -80°C for up to a month. For fewer than 100k cells, the volume of RLT was reduced to 75 µl. The cell lysate was then processed following the manufacturer’s protocol for RNA extraction. The quality of total RNA was checked on the 4200 TapeStation System (Agilent). The SMART-Seq Ultra Low Input RNA Kit for Sequencing (Takara, Cat No.: 634890) was used for the cDNA library construction following the manufacturer’s instructions. The quality and size distribution of the library were checked with an Agilent 5400 bioanalyzer, and the concentration of cDNA was measured by Qubit Fluorometer (Invitrogen). Quantified libraries will be pooled and sequenced on Illumina NovaSeq 6000. The library preparation and sequencing of the marker lines *AtML1*, *AtHB8*, *PIN1*, *SHR*, and *STM* were finished by Novogene Company Limited (Cambridge, United Kingdom), while the maker lines AS2, CO2, FIL, and KN were sent to DRESDEN-concept Genome Center (Dresden, Germany) for library preparation and sequencing.

### snRNA-seq

The construction of the snRNA-seq library was performed using the Chromium Single Cell 3ʹ Version 3 Reagent Kits (10xGemonics). Approximately 10,000 nuclei were processed for gel bead-in-emulsion (GEM) generation utilizing the Chromium Controller. Within these GEMs, the polyadenylated mRNA underwent reverse transcription to synthesize cDNA. Following cDNA amplification and purification, the library was assembled in alignment with the instructions provided in the kit manual. The library fragment size was analyzed using the Agilent 4200 Tapestaion system, and its concentration was measured with the Qubit Fluorometer. The qualified snRNA-seq library was sent to DRESDEN-concept Genome Center (Dresden, Germany), where it underwent sequencing on the Illumina NovaSeq platform.

### Chip-seq

ChIP-seq experiments were conducted with slight modifications to the protocol outlined in the referenced publication^93^. The inflorescence from pro*MYB16*-g*MYB16*: GFP which does not exceed stage 13 (unopened buds) was collected on ice. These samples underwent fixation in MC buffer containing 1% formaldehyde, with a 30-minute vacuum treatment to enhance solution permeation into the tissue. The cross-linked inflorescence was used for chromatin immunoprecipitation experiments with GFP-antibody (Abcam, Cat No.: ab290). The protein-DNA complex was captured by protein A Dynabeads (Thermo Fisher Scientific, Cat No.:10001D), and the subsequent steps involved purifying the DNA and perpetrating the library. The sequencing was carried out on the Illumina NovaSeq platform. This experiment included two biological replicates alongside matching input samples serving as controls.

### Immunoprecipitation-Mass Spectrometry

Protein immunoprecipitation and mass spectrometry followed a previously established protocol^94^. Samples were collected 8 days after bolting, specifically selecting inflorescences younger than stage 13 (unopened buds). Approximately 0.5 grams of pro*MYB16*:*MYB16*-GFP plant material was ground into a fine powder using a pestle and mortar under liquid nitrogen.

The ground tissue was transferred to a 50 ml tube for nuclei isolation. The powdered tissue was resuspended in 20 ml of M1 buffer (10 mM sodium phosphate buffer, 0.1 M NaCl, 1 M 2-methyl 2,4-pentanediol, 10 mM β-mercaptoethanol, and complete protease inhibitor cocktail) and filtered through a 55 µm mesh cloth. The filtrate was centrifuged at 1,000 g for 20 minutes at 4°C to pellet the nuclei. The nuclei pellet was washed with M2 buffer (10 mM sodium phosphate buffer, 0.1 M NaCl, 1 M 2-methyl 2,4-pentanediol, 10 mM β-mercaptoethanol, 10 mM MgCl2, 0.5% Triton X-100, and complete protease inhibitor cocktail) and M3 buffer (10 mM sodium phosphate buffer, 0.1 M NaCl, 10 mM β-mercaptoethanol, and complete protease inhibitor cocktail), and then resuspended in 1 ml of lysis buffer from the μMACS GFP Isolation Kit (Miltenyi Biotec, Cat No.: 130-091-288).

The nuclear suspension was supplemented with protease inhibitor and Benzonase nuclease and sonicated using a Covaris S220 sonicator (peak power = 140.0, cycles/burst = 200, duty factor = 10.0, duration = 180 sec; temp = 6°C). The lysate was centrifuged at 20,000 g for 10 minutes at 4°C twice to remove debris. The supernatant containing soluble nuclear proteins was transferred to a new tube and incubated with 50 µl of anti-GFP MicroBeads for 1 hour at 4°C with gentle rotation. The mixture was then applied to a μ Column (Miltenyi Biotec, Cat No.: 130-042-701) placed in a μMACS Separator. The column was washed six times with 200 µl of lysis buffer and twice with 200 µl of Wash Buffer 2. Proteins were eluted with 50 µl of freshly prepared 8 M urea.

To reduce the cysteine residues in the samples, 7.5 mM DTT was added to the IP eluate, followed by incubation for 30 minutes at 56°C. Alkylation was then performed with 15 mM iodoacetamide (IAA) for 20 minutes in the dark at room temperature.

Proteins were digested overnight at 37°C with 0.5 µg of trypsin (Trypsin Gold, Mass Spectrometry Grade; Promega). The resulting peptide mixture was desalted using Oasis HLB µElution columns (Waters) and dried using a SpeedVac before MS analysis.

The dried peptides were resuspended in 25 µL of 0.1% formic acid/2% acetonitrile (ACN) before measurements. The resuspended peptides were then vortexed and sonicated in a water-bath sonicator for 10 minutes. The peptide concentration was then measured using the NanoDrop One A205 method. Transfer the peptide solution to an autosampler vial, cap it, and place it in the autosampler. Start the “Isocratic Flow” script on the Easy nLC1200 system with 5% Solvent B and a flow rate of 300 nL/min. The mass spectrometer (MS) was set to the “ON” state using the Tune software, and a stable ion spray with Total Ion Current Variation less than 10% was ensured. A method was created in the Xcalibur software’s “Instrument Setup” based on the specified parameters mentioned in the reference (Smaczniak, 2023). The “Sequence Setup” in the Xcalibur software was then used to create a run sequence. The analysis was initiated, and data were automatically saved as .raw files.

The acquired raw MS data were processed using Perseus software, following the procedure in the reference^95^. Raw files were imported, and the data was filtered to remove contaminants, reverse hits, and proteins identified only by site. Proteins identified with at least two unique peptides were considered for further analysis.

Label-free quantification (LFQ) values were log2-transformed, and missing values were imputed using the normal distribution to simulate low-abundance protein detection limits. Statistical significance of protein enrichment was assessed using Student’s t-test, with the false discovery rate (FDR) controlled by the Benjamini-Hochberg procedure. Proteins with FDR < 0.05 were considered significantly enriched.

#### Immunofluorescence for MYB16-GFP and DRMY1-mCherry Colocalization

Inflorescences of transgenic MYB16-GFP and DRMY1-mCherry plants were dissected to reveal early stages of floral organogenesis. The tissues were then embedded in pre-chilled OCT and frozen in an isopentane bath on dry ice.

Cryosections of 10 μm thickness were prepared with a cryotome at −20 °C and collected on poly-lysine coated microscopy slides. The sections were fixed with 4% paraformaldehyde (PFA) in PBS (pH 7.4) for 30 minutes at room temperature and washed three times with 1× PBS. To quench residual aldehyde groups, sections were washed three times with 20 mM glycine in PBS for 10 minutes each.

Permeabilization was performed by incubating the sections in 1% Triton X-100 in PBS for 5 minutes, followed by three washes with PBS. 8% BSA blocking solution prepared in PBS was added to the sections for 1.5 hours at room temperature to minimize non-specific antibody binding. Sections were then incubated overnight at 4 °C with the following primary antibodies diluted in blocking buffer: anti-GFP (1:200, ChromoTek, gb2AF488) and anti-mCherry (1:50, Thermo Fisher, AB_2536611).

The next day, the sections were washed three times with PBS and with the same 8% BSA blocking solution for 10 minutes three times. Subsequently, they were incubated with the secondary antibody (1:500, goat anti-rat IgG H&L Alexa Fluor 555, Thermo Fisher, AB_2535855) in the dark for 1.5 hours at room temperature.

After three final washes with PBS, the sections were counterstained with DAPI for 5 minutes, washed briefly in PBS, and mounted for imaging.

Immunofluorescence acquisitions were taken with a Leica Stellaris8 confocal microscope using a 63x oil immersion objective. Zstacks acquisitions of DAPI-stained nuclei, MYB16-GFP, and DRMY1-mCherry positive nuclei were taken using the 405nm, 488nm, and 555nm lasers respectively followed by lightning deconvolution. Nuclear slices were manually segmented followed by Otsu thresholding. To determine protein colocalisation, the Pearson correlation was calculated followed by the Manders’ overlap coefficients M1 and M2. Visualisation of the nuclei was done with ImageJ95.

#### Scanning Electron Microscopy (SEM)

To examine epidermal surface morphology, scanning electron microscopy (SEM) was performed on wild-type (WT), myb16, myb106, myb16 myb106 double mutant, proMYB16:MYB24-CDS-GFP, and proMYB16:MYB26-CDS-GFP plants. Samples were fixed and prepared following the methanol fixation method as described by Talbot and White ^96^ to improve tissue preservation.

Fresh floral tissues were fixed in 100% methanol for 20 minutes, with vacuum infiltration applied if the tissue did not sink. Fixed samples were transferred to 100% dry ethanol for 30 minutes and further dehydrated in fresh 100% dry ethanol for an additional 30 minutes.

Following dehydration, samples were subjected to critical point drying according to the manufacturer’s instructions. Dried tissues were then mounted onto aluminum SEM stubs using conductive carbon adhesive tabs. Samples were coated with a thin layer of gold using a sputter coater for high-vacuum SEM. Imaging was performed using a Scanning Electron Microscope (LEO Electron Microscopy, Zeiss).All samples were stored in a low-humidity environment (controlled humidity cabinet) before and after SEM imaging to prevent artifacts due to moisture absorption.

Quantification of epidermal cell size was performed using ImageJ. SEM images of the abaxial petal surface were used to measure the length of individual epidermal cells. For each genotype, at least 50 cells were measured from multiple biological replicates. The scale bar embedded in the SEM image was used to calibrate measurements, and the straight-line tool in ImageJ was applied to determine cell length along the longest axis. Measurements were exported to Microsoft Excel and analyzed for statistical significance. Statistical comparisons between genotypes were performed using one-way ANOVA followed by Tukey’s post hoc test, with p-values < 0.05 considered significant.

### Bioinformatics analysis

#### RNA-seq Analysis

First, the quality of the raw sequencing files was evaluated with FastQC (v.0.11.9)^97^. Next, reads were mapped to the A. thaliana genome (TAIR 10) and the Araport11 gene annotation using STAR (v.2.7.9a) with default parameters^98^. Subsequently, samtools (v.1.13)^99^ was used to convert aligned reads into “.bam” format, sort reads by coordinate, remove reads failing platform quality checks and remove reads with a mapping quality < 40. To visualize the coverage of aligned reads, the command “bamCoverage’’ of the deeptools package (v.3.5.1)^100^ was used with the parameters bs 1, - - skipNonCoveredRegions, - - normalizeUsing CPM. In order to obtain read count matrices of mapped reads, the tool “featureCounts” (v.2.0.2)^101^ with default parameters was used. Differential gene expression analysis was performed with functions of the DEseq2 (v.1.46.0)^102^ package separately for sequencing data of the time resolved tissue gene expression atlas and the gene expression data for the myb-family mutants. In both cases raw counts were normalized using the VST function of the DEseq2 package.

#### snRNA-seq Analysis

The .fastq files were processed with Cell Ranger (v.9.0.0) using default parameter values with the A. thaliana genome assembly TAIR 10 and the Araport11 gene annotation. Processing of the raw count matrices included the removal of genes encoded in the organelles, removal of cells that express less than 300 genes and removal of genes that are expressed in less than 10 cells. Next, dataset integration, read count normalization and clustering were done with the R package Seurat (v.5.1.0)^103^. Specifically, all dataset was first normalized and scaled, variable features were detected and Principal Component Analysis (PCA) was applied separately. Then datasets were integrated based on the embedding in PCA space using Canonical Correlation analysis (“CCAIntegration”) with the command IntegrateLayers. This was done for two different combinations of datasets. For the first integration task, all datasets (Ler mature flower, ecotypes and epidermal cells) were combined generating a big atlas containing a mixture of tissues and cell types from different developmental stages. In the second integration task, only single-nuclei datasets from epidermal cells were integrated.

In both cases, a UMAP embedding was computed with the function RunUMAP() based on the embedding generated by the “CCAIntegration” approach from Seurat with the parameters “dims = 1:15” and “n.components = 2”. Next, a neighborhood graph was constructed via the function FindNeighbors() within the space generated by the “CCAIntegration” approach using the parameters “reduction = ‘integrated.cca’” and “dims = 1:15”. In the next step, clustering was performed on that graph with the function FindCluster() setting the resolution parameter to 0.8. In the last step, marker genes for each cluster were identified with the function FindAllMarkers() using the parameter values “only.pos = TRUE”, slot = “data”, min.pct = 0.05, logfc.threshold = 2.

In order to annotate the identified clusters, the average expression in each cluster for a set of genes based on the intersection of the top 100 cluster marker genes and marker genes specific for different tissue types, cell types and developmental time points was used. The sequencing data used for the generation of marker genes for various organ/tissue-types and developmental time points that have not been generated within this study are publicly available with the accession numbers PRJNA79729, PRJNA314076, PRJNA471232 and PRJNA595605.

#### Pseudotime Prediction

The prediction of RNA velocity based on the difference in spliced and unspliced transcripts was performed with velocyto (v.0.17.17)^104^. The detailed modeling of RNA velocities to estimate transcriptional dynamics based on the output of velocyto was conducted with the Python package scvelo (v.0.3.2)^105^. In particular, the scvelo package was used to generate an RNA velocity visualization in UMAP space in the form of an embedding stream. Furthermore, to predict a pseudo-temporal progression of cell states, start and end points in the UMAP embedding for different annotated clusters were selected manually informed by the predicted velocity stream on the entire UMAP embedding. For the downstream analysis of manually defined trajectories of cell states, cells were binned into pseudo time windows. The validity of the predicted pseudo time progression was ensured by comparing the transcriptomes of respective cells in various pseudo time bins to developmental time resolved bulkRNA-seq data.

#### ChIP-seq data processing and peak calling

Raw sequencing reads were trimmed to remove adapter sequences using Trimmomatic(v0.39)^106^, and data quality was assessed with **FastQC** (v0.12.1). Cleaned reads were aligned to the *Arabidopsis thaliana* TAIR10 reference genome using **Bowtie2** (v2.5.1)^107^ with default parameters. Subsequently, unmapped reads, secondary alignments, reads failing platform/vendor quality checks, PCR or optical duplicates, and reads with a mapping quality (MAPQ) score below 40 were filtered out using **Picard MarkDuplicates** (v2.27.5) and **Samtools** (v1.16.1)^108^. For visualization, normalized genome coverage tracks (RPGC, 1× normalization) were generated using **deepTools** *bamCoverage* (v3.5.5)^109^, and displayed as BigWig files. Peaks were called using **MACS2** (v2.2.7.1)^110^, comparing ChIP samples against matched Input controls.

#### Binding Motif Prediction

For the generation of position weight matrices that represent binding motifs, sequences of the top 500 ChIP-seq binding sites, based on peak score, were collected (+/- 100 bp around peak summit) and compared to a set of 500 randomly samples sequences across the genome using functions of the R package memes (v.1.14.0)^111^. Specifically, for the de novo prediction of position weight matrices the command runMeme() with the parameters “objfun=de”, “mod=zoops”, “minw=12”, “maxw=16”, “p=15” and “nmotifs=12” was used. For the comparative analysis using known position weight matrices of already characterized transcription factors, the command runAME() with default parameters and position weight matrices from the DAP-seq based database “ArabidopsisDAPv1.meme” was used.

#### Gene Expression Network analysis

The prediction of gene co-expression networks was performed with the R package GENIE3 (v.1.27.0)^112^ with the parameter “nTrees=100” and otherwise default parameters. Prior to running GENIE3, the expression matrix for a particular group (cluster or pseudo time point) was filtered to only contain genes that are expressed in at least 1/5 of the respective cell population. In order to obtain comparable predicted interaction networks, it was ensured that all batch runs of GENIE3 contained the same genes even if they were previously marked as not being expressed in enough cells.

#### GO-Term Analysis

For the analysis of enriched GO-Terms within certain groups of genes based on clusters or pseudo time bins, the R package topGO (v.2.58.0)^113^ with the parameters “nodeSize=0”, “ontology=BP”, “algor=classic” and “testStatistic=Fisher_Exact” was used. The selected list of genes subjected to GO Term analysis was compared to a background gene list which comprised all remaining genes based on the used annotation (.gtf) file.

## Data availability

The sequencing data generated in this study have been deposited in the NCBI Gene Expression Omnibus (GEO) under accession numbers GSE300242 (ChIP-seq of MYB16), GSE300245 (tissue-specific transcriptomic atlas), and GSE300246 (snRNA-seq).

## Acknowledgements

We thank Peer Martin and the Molekulare Parasitologie group at Humboldt University for assistance with scanning electron microscopy (SEM). Siye Chen was supported by a scholarship from the China Scholarship Council (CSC, Grant No. 202007720063). This work was supported by a DFG project (project number 438774542). Kerstin Kaufmann wishes to thank DFG for funding (458750707, 546593285, 428987924, 512328399, 512328399).

## Author contributions

S.C., M.N., and K.K. jointly conceived and designed the study. M.N. and S.C. jointly developed the analysis strategy, analyzed the experimental data, and contributed to writing the manuscript. S.C. performed the single-cell, microscopy, and molecular experiments, and wrote the manuscript. Z.H. and X.Z. constructed the interactive web portal. F.C. conducted the immunofluorescence experiments and analyzed the corresponding data. C.S. analyzed the ChIP-seq data and contributed to the analysis of IP–MS experiments. C.B. and S.C. performed FANS experiments. S.R. analyzed the phenotypes of MYB mutant lines. Y.Z. quantified cell lengths in promoter-swap lines. K.K. supervised the study and provided input on experimental design and manuscript preparation.

## Competing interests

The authors declare no competing interests.

## Notes

### Competing Interest Statement

The authors have declared no competing interest.

## Reference list

1 Lopez-Anido, C. B. et al. Single-cell resolution of lineage trajectories in the Arabidopsis stomatal lineage and developing leaf. Dev Cell 56, 1043–1055 e1044 (2021). 10.1016/j.devcel.2021.03.014

2 Ryu, K. H., Huang, L., Kang, H. M. & Schiefelbein, J. Single-Cell RNA Sequencing Resolves Molecular Relationships Among Individual Plant Cells. Plant Physiol 179, 1444–1456 (2019). 10.1104/pp.18.01482

3 Denyer, T. et al. Spatiotemporal Developmental Trajectories in the Arabidopsis Root Revealed Using High-Throughput Single-Cell RNA Sequencing. Dev Cell 48, 840–852 e845 (2019). 10.1016/j.devcel.2019.02.022

4 Xu, X., Smaczniak, C., Muino, J. M. & Kaufmann, K. Cell identity specification in plants: lessons from flower development. J Exp Bot 72, 4202–4217 (2021). 10.1093/jxb/erab110

5 Smyth, D. R., Bowman, J. L. & Meyerowitz, E. M. Early flower development in Arabidopsis. Plant Cell 2, 755–767 (1990). 10.1105/tpc.2.8.755

6 Scott, R. J., Spielman, M. & Dickinson, H. G. Stamen structure and function. Plant Cell 16 Suppl, S46–60 (2004). 10.1105/tpc.017012

7 Robinson-Beers, K., Pruitt, R. E. & Gasser, C. S. Ovule Development in Wild-Type Arabidopsis and Two Female-Sterile Mutants. Plant Cell 4, 1237–1249 (1992). 10.1105/tpc.4.10.1237

8 Yang, S. L., Tran, N., Tsai, M. Y. & Ho, C. K. Misregulation of MYB16 expression causes stomatal cluster formation by disrupting polarity during asymmetric cell divisions. Plant Cell 34, 455–476 (2022). 10.1093/plcell/koab260

9 Oshima, Y. et al. MIXTA-like transcription factors and WAX INDUCER1/SHINE1 coordinately regulate cuticle development in Arabidopsis and Torenia fournieri. Plant Cell 25, 1609–1624 (2013). 10.1105/tpc.113.110783

10 Daneva, A., Gao, Z., Van Durme, M. & Nowack, M. K. Functions and Regulation of Programmed Cell Death in Plant Development. Annu Rev Cell Dev Biol 32, 441–468 (2016). 10.1146/annurev-cellbio-111315-124915

11 Zhang, D. et al. The cysteine protease CEP1, a key executor involved in tapetal programmed cell death, regulates pollen development in Arabidopsis. Plant Cell 26, 2939–2961 (2014). 10.1105/tpc.114.127282

12 Higginson, T., Li, S. F. & Parish, R. W. AtMYB103 regulates tapetum and trichome development in Arabidopsis thaliana. Plant J 35, 177–192 (2003). 10.1046/j.1365-313x.2003.01791.x

13. 13 Klepikova, A. V., Logacheva, M. D., Dmitriev, S. E. & Penin, A. A. RNA-seq analysis of an apical meristem time series reveals a critical point in Arabidopsis thaliana flower initiation. BMC Genomics 16, 466 (2015). 10.1186/s12864-015-1688-9

14 Klepikova, A. V., Kasianov, A. S., Gerasimov, E. S., Logacheva, M. D. & Penin, A. A. A high resolution map of the Arabidopsis thaliana developmental transcriptome based on RNA-seq profiling. Plant J 88, 1058–1070 (2016). 10.1111/tpj.13312

15 Ron, M. et al. Hairy root transformation using Agrobacterium rhizogenes as a tool for exploring cell type-specific gene expression and function using tomato as a model. Plant Physiol 166, 455–469 (2014). 10.1104/pp.114.239392

16 Marques-Bueno, M. D. M. et al. A versatile Multisite Gateway-compatible promoter and transgenic line collection for cell type-specific functional genomics in Arabidopsis. Plant J 85, 320–333 (2016). 10.1111/tpj.13099

17 Baima, S. et al. The arabidopsis ATHB-8 HD-zip protein acts as a differentiation-promoting transcription factor of the vascular meristems. Plant Physiol 126, 643–655 (2001). 10.1104/pp.126.2.643

18 Helariutta, Y. et al. The SHORT-ROOT gene controls radial patterning of the Arabidopsis root through radial signaling. Cell 101, 555–567 (2000). 10.1016/s0092-8674(00)80865-x

19 Long, J. A., Moan, E. I., Medford, J. I. & Barton, M. K. A member of the KNOTTED class of homeodomain proteins encoded by the STM gene of Arabidopsis. Nature 379, 66–69 (1996). 10.1038/379066a0

20 Clark, S. E., Jacobsen, S. E., Levin, J. Z. & Meyerowitz, E. M. The CLAVATA and SHOOT MERISTEMLESS loci competitively regulate meristem activity in Arabidopsis. Development 122, 1567–1575 (1996). 10.1242/dev.122.5.1567

21 Lukowitz, W., Mayer, U. & Jurgens, G. Cytokinesis in the Arabidopsis embryo involves the syntaxin-related KNOLLE gene product. Cell 84, 61–71 (1996). 10.1016/s0092-8674(00)80993-9

22 Iida, H., Yoshida, A. & Takada, S. ATML1 activity is restricted to the outermost cells of the embryo through post-transcriptional repressions. Development 146 (2019). 10.1242/dev.169300

23 Lu, P., Porat, R., Nadeau, J. A. & O’Neill, S. D. Identification of a meristem L1 layer-specific gene in Arabidopsis that is expressed during embryonic pattern formation and defines a new class of homeobox genes. Plant Cell 8, 2155–2168 (1996). 10.1105/tpc.8.12.2155

24. 24 Siegfried, K. R. et al. Members of the YABBY gene family specify abaxial cell fate in Arabidopsis. Development 126, 4117–4128 (1999). 10.1242/dev.126.18.4117

25 Semiarti, E. et al. The ASYMMETRIC LEAVES2 gene of Arabidopsis thaliana regulates formation of a symmetric lamina, establishment of venation and repression of meristem-related homeobox genes in leaves. Development 128, 1771–1783 (2001). 10.1242/dev.128.10.1771

26 Xu, L. et al. Novel as1 and as2 defects in leaf adaxial-abaxial polarity reveal the requirement for ASYMMETRIC LEAVES1 and 2 and ERECTA functions in specifying leaf adaxial identity. Development 130, 4097–4107 (2003). 10.1242/dev.00622

27 Heidstra, R., Welch, D. & Scheres, B. Mosaic analyses using marked activation and deletion clones dissect Arabidopsis SCARECROW action in asymmetric cell division. Genes Dev 18, 1964–1969 (2004). 10.1101/gad.305504

28 Galweiler, L. et al. Regulation of polar auxin transport by AtPIN1 in Arabidopsis vascular tissue. Science 282, 2226–2230 (1998). 10.1126/science.282.5397.2226

29 Ecker, J. et al. Genome-Wide Analysis of Gene Expression during Early Arabidopsis Flower Development. PLoS Genetics 2 (2006). 10.1371/journal.pgen.0020117

30 Kim, J. Y. et al. Distinct identities of leaf phloem cells revealed by single cell transcriptomics. Plant Cell 33, 511–530 (2021). 10.1093/plcell/koaa060

31. 31 Shi, D., Lebovka, I., Lopez-Salmeron, V., Sanchez, P. & Greb, T. Bifacial cambium stem cells generate xylem and phloem during radial plant growth. Development 146 (2019). 10.1242/dev.171355

32 Cho, H. et al. Translational control of phloem development by RNA G-quadruplex-JULGI determines plant sink strength. Nat Plants 4, 376–390 (2018). 10.1038/s41477-018-0157-2

33 Etchells, J. P. & Turner, S. R. The PXY-CLE41 receptor ligand pair defines a multifunctional pathway that controls the rate and orientation of vascular cell division. Development 137, 767–774 (2010). 10.1242/dev.044941

34 Crawford, B. C. & Yanofsky, M. F. HALF FILLED promotes reproductive tract development and fertilization efficiency in Arabidopsis thaliana. Development 138, 2999–3009 (2011). 10.1242/dev.067793

35 Mizzotti, C. et al. SEEDSTICK is a master regulator of development and metabolism in the Arabidopsis seed coat. PLoS Genet 10, e1004856 (2014). 10.1371/journal.pgen.1004856

36 Lin, I. W. et al. Nectar secretion requires sucrose phosphate synthases and the sugar transporter SWEET9. Nature 508, 546–549 (2014). 10.1038/nature13082

37 Xu, P. et al. Efficient targeted T-DNA integration for gene activation and male germline-specific gene tagging in Arabidopsis. Plant J 121, e70104 (2025). 10.1111/tpj.70104

38 Program, C. Z. I. C. S. et al. CZ CELLxGENE Discover: a single-cell data platform for scalable exploration, analysis and modeling of aggregated data. Nucleic Acids Res 53, D886–D900 (2025). 10.1093/nar/gkae1142

39 Megill, C. et al. cellxgene: a performant, scalable exploration platform for high dimensional sparse matrices. (2021). 10.1101/2021.04.05.438318

40 Hensel, L. L., Grbic, V., Baumgarten, D. A. & Bleecker, A. B. Developmental and age-related processes that influence the longevity and senescence of photosynthetic tissues in arabidopsis. Plant Cell 5, 553–564 (1993). 10.1105/tpc.5.5.553

41 Skinner, D. J., Dang, T. & Gasser, C. S. The Arabidopsis INNER NO OUTER (INO) gene acts exclusively and quantitatively in regulation of ovule outer integument development. Plant Direct 7, e485 (2023). 10.1002/pld3.485

42 Welch, D. et al. Arabidopsis JACKDAW and MAGPIE zinc finger proteins delimit asymmetric cell division and stabilize tissue boundaries by restricting SHORT-ROOT action. Genes Dev 21, 2196–2204 (2007). 10.1101/gad.440307

43 Lebel-Hardenack, S., Ye, D., Koutnikova, H., Saedler, H. & Grant, S. Conserved expression of a TASSELSEED2 homolog in the tapetum of the dioecious Silene latifolia and Arabidopsis thaliana. The Plant Journal 12, 515–526 (2008). 10.1046/j.1365-313X.1997.d01-4.x

44 Yang, C., Vizcay-Barrena, G., Conner, K. & Wilson, Z. A. MALE STERILITY1 is required for tapetal development and pollen wall biosynthesis. Plant Cell 19, 3530–3548 (2007). 10.1105/tpc.107.054981

45 Deyhle, F., Sarkar, A. K., Tucker, E. J. & Laux, T. WUSCHEL regulates cell differentiation during anther development. Dev Biol 302, 154–159 (2007). 10.1016/j.ydbio.2006.09.013

46 Yang, C. et al. Transcription Factor MYB26 Is Key to Spatial Specificity in Anther Secondary Thickening Formation. Plant Physiol 175, 333–350 (2017). 10.1104/pp.17.00719

47 Mitsuda, N., Seki, M., Shinozaki, K. & Ohme-Takagi, M. The NAC transcription factors NST1 and NST2 of Arabidopsis regulate secondary wall thickenings and are required for anther dehiscence. Plant Cell 17, 2993–3006 (2005). 10.1105/tpc.105.036004

48 Yadav, S., Jalan, K. & Das, S. Comparative Functional Characterization of nst1, nst2, and nst3 in Arabidopsis thaliana Uncovers Previously Unknown Functions in Diverse Developmental Pathways Beyond Secondary Wall Formation. Plant Molecular Biology Reporter 43, 165–179 (2024). 10.1007/s11105-024-01474-1

49 Pinyopich, A. et al. Assessing the redundancy of MADS-box genes during carpel and ovule development. Nature 424, 85–88 (2003). 10.1038/nature01741

50 Ferguson, A. C. et al. Biphasic regulation of the transcription factor ABORTED MICROSPORES (AMS) is essential for tapetum and pollen development in Arabidopsis. New Phytol 213, 778–790 (2017). 10.1111/nph.14200

51 Bowman, J. L., Drews, G. N. & Meyerowitz, E. M. Expression of the Arabidopsis floral homeotic gene AGAMOUS is restricted to specific cell types late in flower development. Plant Cell 3, 749–758 (1991). 10.1105/tpc.3.8.749

52 Ghelli, R. et al. A Newly Identified Flower-Specific Splice Variant of AUXIN RESPONSE FACTOR8 Regulates Stamen Elongation and Endothecium Lignification in Arabidopsis. Plant Cell 30, 620–637 (2018). 10.1105/tpc.17.00840

53. 53 Ginglinger, J. F. et al. Gene coexpression analysis reveals complex metabolism of the monoterpene alcohol linalool in Arabidopsis flowers. Plant Cell 25, 4640–4657 (2013). 10.1105/tpc.113.117382

54 D’Auria, J. C., Pichersky, E., Schaub, A., Hansel, A. & Gershenzon, J. Characterization of a BAHD acyltransferase responsible for producing the green leaf volatile (Z)-3-hexen-1-yl acetate in Arabidopsis thaliana. Plant J 49, 194–207 (2007). 10.1111/j.1365-313X.2006.02946.x

55 McFarlane, H. E., Shin, J. J., Bird, D. A. & Samuels, A. L. Arabidopsis ABCG transporters, which are required for export of diverse cuticular lipids, dimerize in different combinations. Plant Cell 22, 3066–3075 (2010). 10.1105/tpc.110.077974

56 Vanneste, S. et al. Developmental regulation of CYCA2s contributes to tissue-specific proliferation in Arabidopsis. EMBO J 30, 3430–3441 (2011). 10.1038/emboj.2011.240

57. 57 Makkena, S., Lee, E., Sack, F. D. & Lamb, R. S. The R2R3 MYB transcription factors FOUR LIPS and MYB88 regulate female reproductive development. J Exp Bot 63, 5545–5558 (2012). 10.1093/jxb/ers209

58 Song, S. et al. The Jasmonate-ZIM domain proteins interact with the R2R3-MYB transcription factors MYB21 and MYB24 to affect Jasmonate-regulated stamen development in Arabidopsis. Plant Cell 23, 1000–1013 (2011). 10.1105/tpc.111.083089

59 Pajoro, A. et al. Dynamics of chromatin accessibility and gene regulation by MADS-domain transcription factors in flower development. Genome Biol 15, R41 (2014). 10.1186/gb-2014-15-3-r41

60 Wuest, S. E. et al. Molecular basis for the specification of floral organs by APETALA3 and PISTILLATA. Proc Natl Acad Sci U S A 109, 13452–13457 (2012). 10.1073/pnas.1207075109

61 DS, O. M. et al. Control of reproductive floral organ identity specification in Arabidopsis by the C function regulator AGAMOUS. Plant Cell 25, 2482–2503 (2013). 10.1105/tpc.113.113209

62 Yant, L. et al. Orchestration of the floral transition and floral development in Arabidopsis by the bifunctional transcription factor APETALA2. Plant Cell 22, 2156–2170 (2010). 10.1105/tpc.110.075606

63 Brock, M. T. & Weinig, C. Plasticity and environment-specific covariances: an investigation of floral-vegetative and within flower correlations. Evolution 61, 2913–2924 (2007). 10.1111/j.1558-5646.2007.00240.x

64 Rerie, W. G., Feldmann, K. A. & Marks, M. D. The GLABRA2 gene encodes a homeo domain protein required for normal trichome development in Arabidopsis. Genes Dev 8, 1388–1399 (1994). 10.1101/gad.8.12.1388

65 Hubel, A. & Schoffl, F. Arabidopsis heat shock factor: isolation and characterization of the gene and the recombinant protein. Plant Mol Biol 26, 353–362 (1994). 10.1007/BF00039545

66 Kurdyukov, S. et al. The epidermis-specific extracellular BODYGUARD controls cuticle development and morphogenesis in Arabidopsis. Plant Cell 18, 321–339 (2006). 10.1105/tpc.105.036079

67 Shi, J. X. et al. SHINE transcription factors act redundantly to pattern the archetypal surface of Arabidopsis flower organs. PLoS Genet 7, e1001388 (2011). 10.1371/journal.pgen.1001388

68 Sakuma, Y. et al. Functional analysis of an Arabidopsis transcription factor, DREB2A, involved in drought-responsive gene expression. Plant Cell 18, 1292–1309 (2006). 10.1105/tpc.105.035881

69 Lee, B. H. et al. The Arabidopsis thaliana NGATHA transcription factors negatively regulate cell proliferation of lateral organs. Plant Mol Biol 89, 529–538 (2015). 10.1007/s11103-015-0386-y

70 Nicolas, A. et al. De novo stem cell establishment in meristems requires repression of organ boundary cell fate. Plant Cell 34, 4738–4759 (2022). 10.1093/plcell/koac269

71 Powers, J. et al. Next-generation mapping of the salicylic acid signaling hub and transcriptional cascade. Mol Plant 17, 1558–1572 (2024). 10.1016/j.molp.2024.08.008

72 Kijima, S. T. et al. Control of plasma membrane-associated actin polymerization specifies the pattern of the cell wall in xylem vessels. Nat Commun 16, 1921 (2025). 10.1038/s41467-025-56866-y

73 Adamczyk, B. J., Lehti-Shiu, M. D. & Fernandez, D. E. The MADS domain factors AGL15 and AGL18 act redundantly as repressors of the floral transition in Arabidopsis. Plant J 50, 1007–1019 (2007). 10.1111/j.1365-313X.2007.03105.x

74 Meng, Q. et al. Molecular mechanism of interaction between SHORT VEGETATIVE PHASE and APETALA1 in Arabidopsis thaliana. Plant Physiol Biochem 220, 109512 (2025). 10.1016/j.plaphy.2025.109512

75 Serivichyaswat, P. et al. Expression of the floral repressor miRNA156 is positively regulated by the AGAMOUS-like proteins AGL15 and AGL18. Mol Cells 38, 259–266 (2015). 10.14348/molcells.2015.2311

76 Hasson, A. et al. Evolution and diverse roles of the CUP-SHAPED COTYLEDON genes in Arabidopsis leaf development. Plant Cell 23, 54–68 (2011). 10.1105/tpc.110.081448

77 Pineau, E. et al. CYP77B1 a fatty acid epoxygenase specific to flowering plants. Plant Sci 307, 110905 (2021). 10.1016/j.plantsci.2021.110905

78 Tang, J. et al. GDSL lipase occluded stomatal pore 1 is required for wax biosynthesis and stomatal cuticular ledge formation. New Phytol 228, 1880–1896 (2020). 10.1111/nph.16741

79. 79 Duan, H. & Schuler, M. A. Differential expression and evolution of the Arabidopsis CYP86A subfamily. Plant Physiol 137, 1067–1081 (2005). 10.1104/pp.104.055715

80 Li, S. et al. Arabidopsis ACYL-ACTIVATING ENZYME 9 (AAE9) encoding an isobutyl-CoA synthetase is a key factor connecting branched-chain amino acid catabolism with iso-branched wax biosynthesis. New Phytol 233, 2458–2470 (2022). 10.1111/nph.17941

81 Tanaka, T., Tanaka, H., Machida, C., Watanabe, M. & Machida, Y. A new method for rapid visualization of defects in leaf cuticle reveals five intrinsic patterns of surface defects in Arabidopsis. Plant J 37, 139–146 (2004). 10.1046/j.1365-313x.2003.01946.x

82 Cai, Y. S. et al. Arabidopsis AtMSRB5 functions as a salt-stress protector for both Arabidopsis and rice. Front Plant Sci 14, 1072173 (2023). 10.3389/fpls.2023.1072173

83 Zhong, C. et al. Gibberellic Acid-Stimulated Arabidopsis6 Serves as an Integrator of Gibberellin, Abscisic Acid, and Glucose Signaling during Seed Germination in Arabidopsis. Plant Physiol 169, 2288–2303 (2015). 10.1104/pp.15.00858

84 Fowler, T. J., Bernhardt, C. & Tierney, M. L. Characterization and expression of four proline-rich cell wall protein genes in Arabidopsis encoding two distinct subsets of multiple domain proteins. Plant Physiol 121, 1081–1092 (1999). 10.1104/pp.121.4.1081

85 O’Malley, R. C. et al. Cistrome and Epicistrome Features Shape the Regulatory DNA Landscape. Cell 165, 1280–1292 (2016). 10.1016/j.cell.2016.04.038

86 McCarthy, R. L., Zhong, R. & Ye, Z. H. MYB83 is a direct target of SND1 and acts redundantly with MYB46 in the regulation of secondary cell wall biosynthesis in Arabidopsis. Plant Cell Physiol 50, 1950–1964 (2009). 10.1093/pcp/pcp139

87 Liang, Y. K. et al. AtMYB61, an R2R3-MYB transcription factor controlling stomatal aperture in Arabidopsis thaliana. Curr Biol 15, 1201–1206 (2005). 10.1016/j.cub.2005.06.041

88 Wu, P. et al. DRMY1, a Myb-Like Protein, Regulates Cell Expansion and Seed Production in Arabidopsis thaliana. Plant Cell Physiol 60, 285–302 (2019). 10.1093/pcp/pcy207

89 Huang, H. et al. The DELLA proteins interact with MYB21 and MYB24 to regulate filament elongation in Arabidopsis. BMC Plant Biol 20, 64 (2020). 10.1186/s12870-020-2274-0

90 Schumacher, J., Kaufmann, K. & Yan, W. Multiplexed GuideRNA-expression to Efficiently Mutagenize Multiple Loci in Arabidopsis by CRISPR-Cas9. Bio-Protocol 7 (2017). 10.21769/BioProtoc.2166

91. 91 Clough, S. J. & Bent, A. F. Floral dip: a simplified method for Agrobacterium-mediated transformation of Arabidopsis thaliana. Plant J 16, 735–743 (1998). 10.1046/j.1365-313x.1998.00343.x

92 Moreno-Romero, J., Santos-Gonzalez, J., Hennig, L. & Kohler, C. Applying the INTACT method to purify endosperm nuclei and to generate parental-specific epigenome profiles. Nat Protoc 12, 238–254 (2017). 10.1038/nprot.2016.167

93 Kaufmann, K. et al. Chromatin immunoprecipitation (ChIP) of plant transcription factors followed by sequencing (ChIP-SEQ) or hybridization to whole genome arrays (ChIP-CHIP). Nat Protoc 5, 457–472 (2010). 10.1038/nprot.2009.244

94. 94 Smaczniak, C. in Plant gene regulatory networks: Methods and protocols 163–181 (Springer, 2023).

95 Smaczniak, C. Immunoprecipitation-Mass Spectrometry (IP-MS) of Protein-Protein Interactions of Nuclear-Localized Plant Proteins. Methods Mol Biol 2698, 163–181 (2023). 10.1007/978-1-0716-3354-0_11

96 Talbot, M. J. & White, R. G. Methanol fixation of plant tissue for Scanning Electron Microscopy improves preservation of tissue morphology and dimensions. Plant Methods 9, 36 (2013). 10.1186/1746-4811-9-36

97 Andrews, S. FastQC: a quality control tool for high throughput sequence data., <https://www.bioinformatics.babraham.ac.uk/projects/fastqc/> (2010).

98 Dobin, A. et al. STAR: ultrafast universal RNA-seq aligner. Bioinformatics 29, 15–21 (2013). 10.1093/bioinformatics/bts635

99 Danecek, P. et al. Twelve years of SAMtools and BCFtools. Gigascience 10 (2021). 10.1093/gigascience/giab008

100 Ramirez, F. et al. deepTools2: a next generation web server for deep-sequencing data analysis. Nucleic Acids Res 44, W160–165 (2016). 10.1093/nar/gkw257

101 Liao, Y., Smyth, G. K. & Shi, W. featureCounts: an efficient general purpose program for assigning sequence reads to genomic features. Bioinformatics 30, 923–930 (2014). 10.1093/bioinformatics/btt656

102 Love, M. I., Huber, W. & Anders, S. Moderated estimation of fold change and dispersion for RNA-seq data with DESeq2. Genome Biol 15, 550 (2014). 10.1186/s13059-014-0550-8

103 Satija, R., Farrell, J. A., Gennert, D., Schier, A. F. & Regev, A. Spatial reconstruction of single-cell gene expression data. Nat Biotechnol 33, 495–502 (2015). 10.1038/nbt.3192

104 La Manno, G. et al. RNA velocity of single cells. Nature 560, 494–498 (2018). 10.1038/s41586-018-0414-6

105 Bergen, V., Lange, M., Peidli, S., Wolf, F. A. & Theis, F. J. Generalizing RNA velocity to transient cell states through dynamical modeling. Nat Biotechnol 38, 1408–1414 (2020). 10.1038/s41587-020-0591-3

106 Bolger, A. M., Lohse, M. & Usadel, B. Trimmomatic: a flexible trimmer for Illumina sequence data. Bioinformatics 30, 2114–2120 (2014). 10.1093/bioinformatics/btu170

107 Langmead, B. & Salzberg, S. L. Fast gapped-read alignment with Bowtie 2. Nat Methods 9, 357–359 (2012). 10.1038/nmeth.1923

108 Li, H. et al. The Sequence Alignment/Map format and SAMtools. Bioinformatics 25, 2078–2079 (2009). 10.1093/bioinformatics/btp352

109 Ramirez, F., Dundar, F., Diehl, S., Gruning, B. A. & Manke, T. deepTools: a flexible platform for exploring deep-sequencing data. Nucleic Acids Res 42, W187–191 (2014). 10.1093/nar/gku365

110 Zhang, Y. et al. Model-based analysis of ChIP-Seq (MACS). Genome Biol 9, R137 (2008). 10.1186/gb-2008-9-9-r137

111 Bailey, T. L., Johnson, J., Grant, C. E. & Noble, W. S. The MEME Suite. Nucleic Acids Res 43, W39–49 (2015). 10.1093/nar/gkv416

112 Huynh-Thu, V. A., Irrthum, A., Wehenkel, L. & Geurts, P. Inferring regulatory networks from expression data using tree-based methods. PLoS One 5 (2010). 10.1371/journal.pone.0012776

113. 113 topGO: Enrichment Analysis for Gene Ontology v. 2.59.0 (R package, 2024).

